# Discovery of Novel and Selective GPR17 Antagonists as Pharmacological Tools for Developing New Therapeutic Strategies in Diabetes and Obesity

**DOI:** 10.1101/2024.12.04.626849

**Authors:** Hu Zhu, Jason M. Conley, Surendra Karavadhi, Justin E. LaVigne, Val J. Watts, Hongmao Sun, Min Shen, Matthew D. Hall, Hongxia Ren, Samarjit Patnaik

**Affiliations:** Early Translational Branch, Division of Preclinical Innovation, National Center for Advancing Translational Sciences, National Institutes of Health, Rockville, MD 20850, USA; Herman B. Wells Center for Pediatric Research, Department of Pediatrics, National Institutes of Health, Rockville, MD 20850, USA; Center for Diabetes and Metabolic Diseases, National Institutes of Health, Rockville, MD 20850, USA; Stark Neurosciences Research Institute, National Institutes of Health, Rockville, MD 20850, USA; Department of Anatomy, Cell Biology & Physiology, National Institutes of Health, Rockville, MD 20850, USA; Department of Biochemistry & Molecular Biology, National Institutes of Health, Rockville, MD 20850, USA; Department of Pharmacology & Toxicology, Indiana University School of Medicine, Indianapolis, IN 46202, USA; Department of Medicinal Chemistry and Molecular Pharmacology, College of Pharmacy, Purdue University, West Lafayette, IN 47907, USA

**Keywords:** G protein-coupled receptor (GPCR), high-throughput screening (HTS), signal transduction, cyclic AMP (cAMP), calcium, antagonist, selectivity, glucagon-like peptide 1 (GLP-1), diabetes, obesity

## Abstract

G protein coupled receptors (GPCRs) are promising targets for diabetes and obesity therapy due to their roles in metabolism and excellent potential for pharmacological manipulation. We previously reported that *Gpr17* ablation in the brain-gut axis leads to improved metabolic homeostasis, suggesting GPR17 antagonism could be developed for diabetes and obesity treatment. Here, we performed high throughput screening (HTS) and identified two new GPR17 antagonists (compound 978 and 527). Both compounds antagonized downstream Gαi/o, Gαq and β-arrestin signaling with high selectivity for GPR17, but not the closely related purinergic and cysteinyl leukotriene receptors. The molecular mechanisms of antagonism were revealed through Schild analysis, structure-activity relationship (SAR) studies and homology modelling. Compound 978 and its analog (793) attenuated GPR17 signaling and promoted glucagon-like peptide-1 (GLP-1) secretion in enteroendocrine cells. In summary, we identified selective GPR17 antagonists through HTS, which represent promising pharmacological tools for developing new therapeutic strategies in diabetes and obesity.

**Significance:** Our work highlights the therapeutic potential of GPR17 antagonism in the treatment of diabetes and obesity by leveraging its role in metabolic regulation. In previous studies, we have shown that *Gpr17* ablation improves metabolic homeostasis, and here we expanded our research by identifying two novel small molecule antagonists of GPR17 through high-throughput screening. The compounds inhibited multiple downstream signaling pathways of GPR17 with high selectivity over other closely related receptors. Of particular significance, compound 978 and its analogs not only attenuated GPR17 signaling but also increased glucagon-like peptide-1 (GLP-1) secretion, a critical hormone for glucose homeostasis and appetite regulation. These findings shed new light into the molecular mechanisms of GPR17 antagonism and introduce valuable pharmacological tools for further exploration of therapeutic strategies in diabetes and obesity.

## Introduction

Metabolic diseases represent a spectrum of disorders, including diabetes, obesity, hyperlipidemia, and cardiovascular disorders, due to the disrupted metabolic homeostasis in the human body ^1^. G-protein coupled receptors (GPCRs) are a large family of seven-transmembrane-domain receptors and are of particular interest for pharmaceutical intervention ^2,3^. Approximately 30-40% of all FDA approved drugs modulate GPCRs ^4^. GPCRs in the endocrine tissues respond to metabolic hormones, nutrients, metabolites and neurotransmitters, and therefore are promising therapeutic targets for metabolic diseases ^5^. For instance, the GLP-1 receptor has emerged as one of the most successful targets for treating type 2 diabetes mellitus (T2DM) and obesity, and eight GLP-1 receptor agonists have received recent approval in the US market ^6,7^. These new drugs, however, have high cost and adverse effects mainly associated with gastrointestinal discomfort ^6^. Deorphanization efforts over the past two decades have unveiled a growing number of GPCRs as potential therapeutic targets for metabolic diseases ^2,8^. GPR17 is an orphan GPCR we recently identified as a potential target for intervention in diabetes and obesity ^9–12^. We found that GPR17 was an effector of Forkhead box protein O1 (FoxO1) in hypothalamic agouti-related peptide (AgRP) neurons, a pivotal regulator of feeding behavior and energy expenditure ^11^. Loss-of-function of *Gpr17* in AgRP neurons increased central nervous system sensitivity to insulin and leptin, leading to reduced food intake and increased relative energy expenditure in animal models^10^. Furthermore, GPR17 was found to be expressed in the GLP-1 producing enteroendocrine cells (EECs). Inducible knockout of *Gpr17* in gut epithelium led to increased secretion of GLP-1 and improved glucose tolerance ^12^. Beyond these animal studies, our investigations with human GPR17 revealed distinct downstream signaling profiles associated with *GPR17* missense variants in patients with metabolic diseases, including diabetes and obesity ^13^.

GPR17 was initially identified from a human hippocampus cDNA library using P2Y-homologous sequences ^14^. In the human genome, *GPR17* is located on chromosome 2q21 containing 4 exons. There are four *GPR17* transcript variants, which give rise to two distinct protein isoforms: the long isoform (hGPR17L) and the short isoform (hGPR17S), distinguished by an additional 28 amino acids at the N-terminus of the long isoform ^15^. In contrast, rodent *Gpr17* possesses only one protein isoform, which is an ortholog of hGPR17S ^15^. GPR17 is phylogenetically located at an intermediate position between the purinergic type 2 receptors, subtype Y (P2RYs) and cysteinyl leukotriene receptors (CYSLTRs). Both cysteinyl leukotrienes and uracil nucleotides were reported to activate GPR17, leading to both inhibition of cAMP production and calcium signaling ^16^. However, the results were challenged by other groups ^17–20^.

In addition to the controversy and uncertainty of endogenous ligands of GPR17, the currently available chemical probes in literature have limitations in selectivity and/or potency. For example, 3-(2-carboxyethyl)-4,6-dichloro-1H-indole-2-carboxylic acid (MDL29,951, MDL) was identified as a small molecule agonist ^21^, which was subsequently confirmed by multiple groups including us ^12,13,22^. MDL was initially identified as an antagonist of N-methyl-D-aspartate (NMDA) receptors with high affinity for its glycine binding site (Ki = 140 nM) ^23^. Its activity was confirmed both *in vitro* and *in vivo* ^23,24^. After the discovery of MDL, selective antagonists of CYSLTRs including HAMI3379 (HAMI) ^25^, pranlukast ^21^, and montelukast ^16^ were found to weakly antagonize the activity of GPR17. More GPR17 antagonists with different scaffolds were discovered and published in recent years ^22,26–28^. In our efforts to identify novel small molecule antagonists of GPR17, we developed high-throughput screening (HTS) assays, which were amenable to 1536-well plate format, and screened a collection of libraries containing ∼300K small molecule compounds. Two novel compounds, namely 978 and 527, were successfully identified and validated through a battery of primary assays, counter assays, and orthogonal assays. Both compounds exhibited high selectivity for GPR17, remaining inert to P2RYs and CYSLTRs, and effectively blocked the activation of MDL-induced downstream signaling pathways of GPR17. We revealed the molecular mechanisms of antagonism through Schild analysis, structure-activity relationship (SAR) studies, and homology modelling. Furthermore, our functional studies in enteroendocrine cells showed that the newly identified GPR17 antagonist and the analogs derived from SAR studies attenuated GPR17-mediated inhibition of cAMP signaling and GLP-1 secretion.

## Results

### Assay design, optimization, and miniaturization for HTS

GPR17 couples to Gαi/o, Gαs, Gαq, and β-arrestin signaling pathways ^21^. β-arrestin recruitment after receptor activation was shown to be mediated via both G protein-dependent and independent mechanisms ^21,29^. To identify small molecule antagonists independent of signaling pathways, the PathHunter™ β-arrestin recruitment assay (DiscoveRx/Eurofins) was chosen as the primary assay for our high-throughput screening campaign. In this technology, hGPR17L (GenBank accession number NM_005291) tagged with ProLink™ was stably expressed in a human bone osteosarcoma epithelial (U2OS) cell line which co-expressed β-arrestin 2 protein tagged with the larger enzyme acceptor fragment of β-galactosidase. Ligand-induced receptor activation and subsequent recruitment of β-arrestin to the receptor leads to the fragment complementation of β-galactosidase, which can be monitored by chemiluminescent signal. It is a simple and robust assay with high specificity, and it is compatible for HTS ^19^. MDL, a surrogate agonist of GPR17, was used in our experiments to activate the receptor. In the PathHunter GPR17 β-arrestin recruitment assay, MDL activated GPR17 at EC_50_=0.34 µM which was comparable to previous studies ^21^ (Figure 1A). HAMI, which was initially developed as a cysteinyl leukotriene receptor 2 (CysLTR2) antagonist (IC_50_=3.8 nM) ^20^, was also found to be an antagonist of GPR17 exhibiting an IC_50_ in the single digit micromolar range ^25^. It was used as a reference antagonist compound in our HTS campaign. Pretreatment of PathHunter GPR17 β-arrestin cells with HAMI effectively blocked the activation of receptors induced by an EC_80_ concentration of MDL (2 µM) in a dose dependent manner with an IC_50_ of 8.2 µM (Figure 1B). This IC_50_ for HAMI was also comparable with previous studies ^25^.

**Figure 1.**
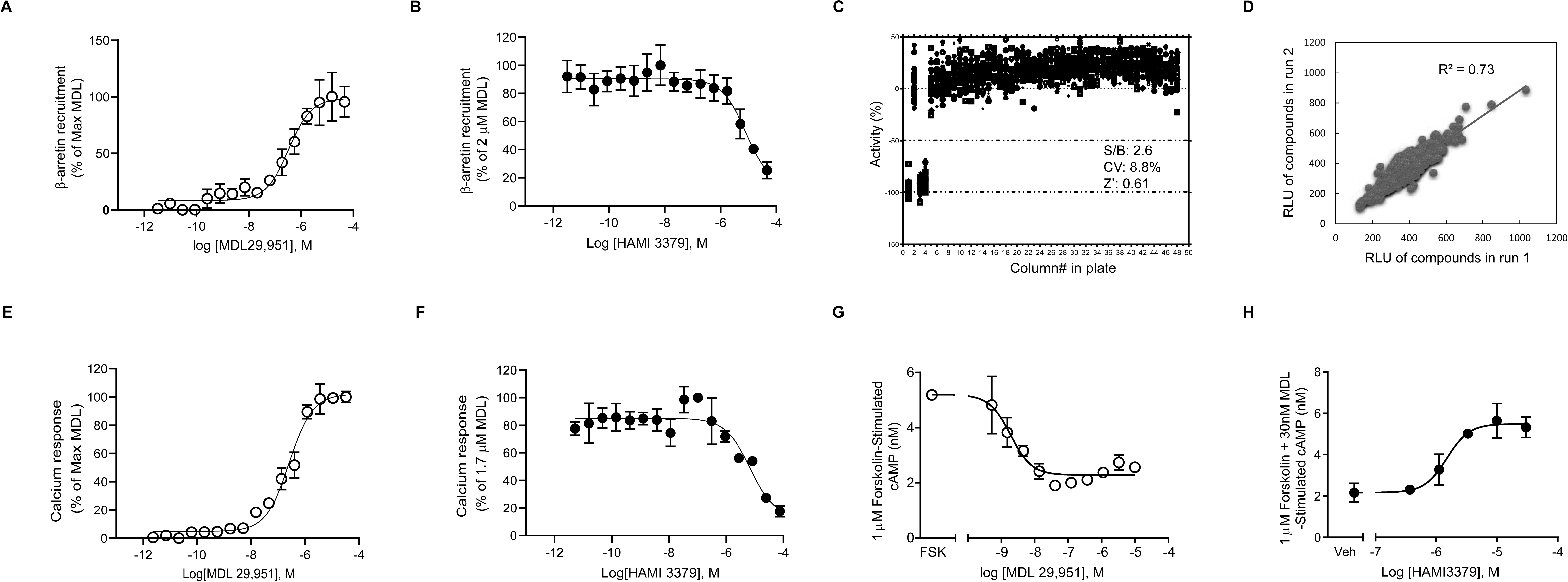
Development of GPR17 assays for high-throughput screening. PathHunter GPR17 β-Arrestin assay was developed and optimized in 1536-well plate format and used as the primary assay for high-throughput screening. **A.** Dose-response curve of MDL compound in agonist mode, EC_50_ of MDL was 0.37µM. **B.** Dose-response curve of HAMI compound in antagonist mode, 2µM MDL (∼EC_80_ concentration) was used to stimulate the receptor. EC_50_ of HAMI was 8.2µM. **C.** Scatter plot of PathHunter GPR17 β-Arrestin assay in a 1536-well plate format. Column 1 was unstimulated control (no MDL). Column 2 was stimulated neural control (2 µM MDL). Columns 3 and 4 were the antagonist control (30 µM HAMI and 2 µM MDL). Column 5 to 48 were pretreated with small molecule library compounds and then stimulated with 2 µM MDL. **D.** Reproducibility test of PathHunter GPR17 β-arrestin recruitment assay from two runs. Calcium mobilization assay was optimized in 1536-well plate format and used as an orthogonal assay. **E.** Dose response curve of MDL compound in the agonist mode. EC_50_ of MDL was 0.28µM. **F.** Dose response curve of HAMI compound in antagonist mode. 1.7µM MDL (∼EC_80_ concentration) was used to stimulate the receptor. EC_50_ of HAMI was 8.1µM. cAMP accumulation assay (Cisbio) was developed in 384-well plate format as a second orthogonal assay. **G**. Dose response curve of MDL compound in agonist mode. EC_50_ of MDL was 1.9 nM. 1 µM forskolin was used to stimulate the production of cAMP. **H**. cAMP accumulation assay in antagonist mode. An increased concentration of HAMI was used in this experiment. 1 µM forskolin was used to stimulate the production of cAMP and 30 nM MDL was used to activate the receptor.

To enable the automated screening of small molecule libraries, the assay was miniaturized to, optimized, and validated in a 1536-well microplate format. The first four columns of the 1536-well plates were reserved as the control columns. Column 1 was unstimulated control (32 wells, no MDL). Column 2 was an agonist stimulated control (32 wells, 2 µM MDL). Columns 3 and 4 were the antagonist control (32 wells each, 30 µM HAMI and 2 µM MDL). Columns 5 to 48 were pretreated with small molecule library compounds and then stimulated with 2 µM MDL. Figure 1C showed an example scatter plot of receptor activity (%) from a 1536-well DMSO control plate to validate the assay. The ratio of signal/background (S/B) was 2.6, coefficient of variation (CV) was 8.8%, and Z factor was 0.61. To assess the reproducibility, we tested a small library of 1,408 compounds in this β-arrestin recruitment assay in two runs. The data showed good reproducibility between both runs in our experiment (R^2^=0.73, Figure 1D).

To further validate the hits identified from HTS, we also developed calcium mobilization and cAMP accumulation assays as orthogonal assays using a 1321N1 cell line (human astrocytoma cell line) that stably expresses hGPR17L. A Calbryte-520 probenecid-free and wash-free calcium assay was used to detect calcium mobilization. Cells were treated with increasing concentrations of MDL and calcium influx was measured by a Fluorometric Imaging Plate Reader (FLIPR) Penta high-throughput cellular screening system. MDL activated the receptor at EC_50_=0.28 µM (Figure 1E), and HAMI effectively blocked the receptor activation stimulated by MDL (at EC_80_ concentration of 1.7 µM) at IC_50_=8.1 µM (Figure 1F). GPR17-mediated cAMP signaling was measured with a homogeneous time resolved fluorescence (HTRF) cAMP assay.

Cells were treated with varying concentrations of MDL, and subsequently stimulated with the direct adenylyl cyclase activator forskolin (1 µM). MDL activated GPR17 at EC_50_=1.9 nM (Figure 1G), and HAMI effectively blocked the receptor activation by MDL (at EC_80_ concentration of 30 nM) at IC_50_=1.5 µM (Figure 1H).

### Identification and validation of antagonists of GPR17 from HTS

To identify antagonists of GPR17, we conducted a high-throughput screening campaign following a rigorous and stringent process (Figure 2A). We screened in-house libraries consisting of a total number of 301,408 small molecule compounds in 1536-well format using the PathHunter GPR17 β-arrestin recruitment assay as the primary assay. The libraries were screened at a final concentration of compounds at 66.7 µM. Briefly, the cells were pretreated with the library compounds for 10 min and then stimulated with 2 µM MDL (EC_80_ concentration). The activity of each compound was normalized to 66.7 µM HAMI as the positive control (−100%) and DMSO as the neutral control (0%). The activity of a compound equal or less than −50% was designated as a hit. According to this criteria, 1,378 hits were identified and further assessed in the PathHunter GPR17 β-arrestin recruitment assay at 11 doses (a series of 5-fold dilutions from 66.7 µM to 0.0068 nM). 380 hits were successfully confirmed, and the confirmation rate was 27.6% (Figure 2A). Figure 2B and 2C shows the structures and dose response curves of two confirmed hit compounds, namely compound 978 and 572.

**Figure 2.**
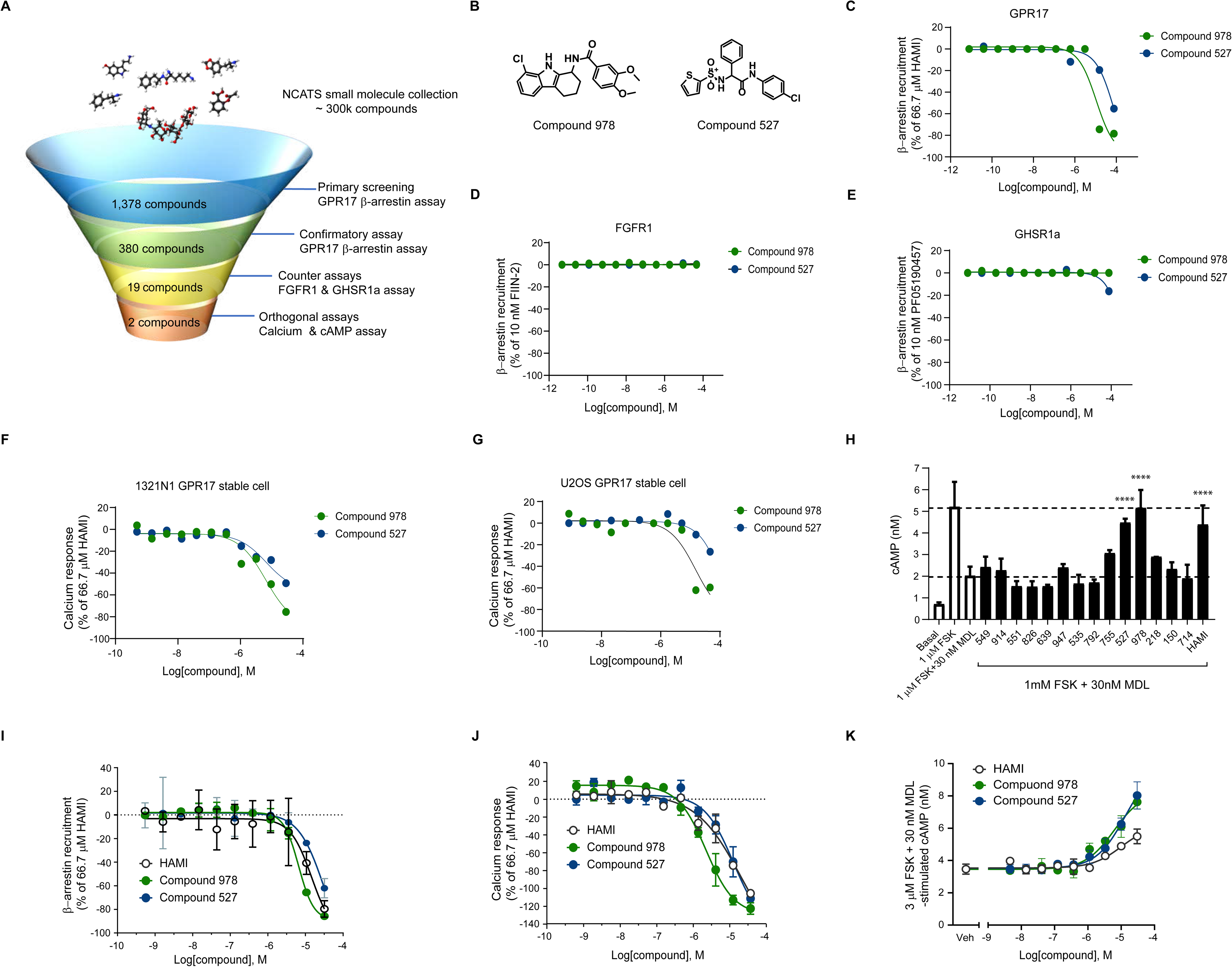
Identification of antagonists of GPR17 by high-throughput screening. **A.** Flow chart of high-throughput screening. PathHunter GPR17 β-arrestin assay was used as the primary assay for high-throughput screening campaign. 301,408 small molecule compounds from in-house libraries were screened at single concentration (66.7 µM) by the primary assay. 1,378 compounds, of which the max response was more than 50%, were cherry-picked as hits and tested in 11 doses in the PathHunter GPR17 β-arrestin assay. The activity of 380 compounds was confirmed by confirmatory assay. The artifacts of PathHunter assay were triaged by two counter assays, PathHunter FGFR1 functional assay and PathHunter GHSR1α β-arrestin assay. 19 of 380 hits were inactive in two counter assays and were selected for further validation in two orthogonal assays, calcium mobilization assay and cAMP accumulation assay. Two antagonists (compounds 978 and 527) were finally validated in these two orthogonal assays. **B.** Structure of compounds 978 and 527. **C.** Dose response curve of compounds 978 and 527 in PathHunter U2OS GPR17 β-arrestin assay. **D.** Dose response curve of compounds 978 and 527 in PathHunter U2OS FGFR1 functional assay. **E.** Dose response curve of compounds 978 and 527 in PathHunter U2OS GHSR1α β-arrestin assay. **F.** Dose response curve of compounds 978 and 527 in calcium mobilization assay (1321N1 GPR17 stable cell line). **G.** Dose response curve of compounds 978 and 527 in calcium mobilization assay (U2OS GPR17 β-arrestin cell line). **H.** Attenuation of MDL-induced inhibition of forskolin-stimulated cAMP accumulation in 1321N1 GPR17 stable cells. Cells were pretreated with 30 µM of compounds and then stimulated with 1 µM forskolin and 30 nM MDL. The date was analyzed by one-way ANOVA with Dunnett’s posttest. Compound 978 was re-purchased and subjected to purification, and compound 527 was re-synthesized by our chemist. The purified compounds were reassessed in three GPR17 assays, PathHunter GPR17 U2OS β-arrestin assay (**I**), calcium mobilization assay (1321N1 GPR17 stable cell line) (**J**), cAMP HTRF assay (1321N1 GPR17 stable cell line) (**K**). HAMI was used as a reference compound. The IC_50_ of 978 was 6.6 µM (arrestin assay), 2.3 µM (calcium mobilization assay) and 6.1 µM (cAMP HTRF assay). The IC_50_ of 527 was 33.3 µM (arrestin assay), 13 µM (calcium mobilization assay) and 13.2 µM (cAMP HTRF assay).

To triage the assay artifacts and nonselective hits, two other PathHunter assays were chosen as the counter assays. Fibroblast growth factor receptor 1 (FGFR1) is a receptor tyrosine kinase (RTK), and activation of receptor leads to the recruitment of downstream SH2 domain proteins, which was engineered in the PathHunter β-galactosidase complementation assay for detection of receptor activation. The FGFR1 functional assay was used as a counter assay to triage interference compounds that modulate FGFR1 and β-galactosidase. Growth hormone secretagogue receptor 1a (GHSR1a) is a functional GPCR for gut peptide ghrelin. GHSR1a β-arrestin recruitment assay was used as another counter assay to triage false positive hits that may induce β-arrestin recruitment non-selectively. The performance of counter assays was shown in supplemental figure 1. 19 out of 380 hits (5%) identified from the confirmatory assay were shown to be inactive in both the FGFR1 functional assay and GHSR1a β-arrestin recruitment assay (Figure 2A). The dose response curves of two compounds (978 and 572) showed no activity in the counter assays (Figure 2D&E).

As the primary screen and counter screen were conducted in the PathHunter β-arrestin recruitment assay, activation of G-protein signaling pathways by 19 hits was further evaluated to triage assay artifacts. Activation of the Gαq signaling pathway was evaluated in two GPR17 stable cell lines with different cell backgrounds (1321N1 cell line and U2OS cell line), and antagonism of the Gαi/o signaling pathway by the compounds was evaluated in the 1321N1 GPR17 stable cell line. The different cell background, different readouts, and distinct signaling pathways were expected to further triage artifacts identified from the primary assay. In these orthogonal assays, 2 out of 19 hits, compounds 978 and 572, were confirmed. Compound 978 had a tetrahydro-1H-carbazole core while compound 527 had a sulfonamido acetamide core in a dipeptide-like backbone (Figure 2B). Compounds 978 and 572 dose dependently reduced calcium signaling (Figure 2F&G). Furthermore, 10µM compounds 978 and 572 significantly antagonized the MDL-mediated inhibition of forskolin-stimulated cAMP accumulation (Figure 2H).

Contaminants or decomposition products from library compounds are a source of pan-assay interference compounds (PAINS) ^30^. To exclude this possibility, compounds 978 and 527 were either re-purchased or re-synthesized, subjected to purification, and reassessed in three GPR17 assays. The antagonistic activity of these two compounds was successfully validated in the β-arrestin recruitment assay, calcium mobilization assay, and cAMP accumulation assay. As a reference, HAMI was included in the experiments. The potency of compound 978 was moderately higher than HAMI, while the potency of compound 527 was similar or lower than HAMI (Figure 2I-K). In the β-arrestin recruitment, calcium mobilization and cAMP accumulation assays, the IC_50_ of compound 978 was 6.6 µM, 2.3 µM and 6.1 µM, respectively; the IC_50_ of compound 527 was 33.3 µM, 13 µM and 13.2 µM, respectively; the IC_50_ of HAMI was 14.6 µM, 21 µM and 10 µM, respectively.

### Characterization of antagonists of GPR17

From a phylogenetic point of view, GPR17 is in an intermediate position between the P2RY and CYSLTR families ^31^. Therefore, the receptors from these two families were chosen to further evaluate the selectivity of compounds 978 and 527. First, we assessed the selectivity of compounds at a fixed dose in cells transiently overexpressing hGPR17L, CYSLTR1, CYSLTR2 and P2RY1. As expected, 10 µM of compound 978 and 527 blocked calcium signaling of hGPR17L receptor stimulated by 200 nM MDL (Figure 3A). 10 µM of compound 978 and 527 failed to block the activation of CYSLTR1 (Figure 3B), CYSLTR2 (Figure 3C), and P2RY1 (Figure 3D) stimulated by their respective ligands. In contrast, pranlukast, HAMI, and MRS2179, as the positive controls, antagonized CYSLTR1, CYSLTR2, and P2RY1, respectively. Secondly, we tested the selectivity of 978 and 527 in a dose dependent manner in the stable cell lines. The activation of CYSLTR1, CYSLTR2, and P2RY1 was evaluated by calcium mobilization assay in CYSLTR1, CYSLTR2, and P2RY1 HEK293 stable cell lines and ACTone cAMP assay ^32^ in the P2RY12 HEK293 stable cell line. The CYSLTR1 and CYSLTR2 were activated by their endogenous ligand, leukotriene D4 (LTD4), with an EC_50_ of 3 nM (Supplemental figure 2A) and 4.2 nM (Supplemental figure 2B) respectively. The P2RY1 was activated by its endogenous ligand, adenosine triphosphate (ATP), at EC_50_ of 4.6 nM (Supplemental figure 2C). The EC_50_ of 2-Methylthioadenosine diphosphate (2MeSADP) was 0.032 nM measured by ACTone cAMP assay in the P2RY12 HEK293 stable cell line (Supplemental figure 2D). The antagonist activities of 978 and 527 were evaluated in these stable cell lines by stimulating them with their respective ligands at approximately EC_80_ concentrations. The data showed that 978 exhibited some inhibitory effect (∼ 41%) on activation of CYSLTR1 receptor only at the highest assay concentration of 33 µM (Figure 3E), but not the other three receptors (Figures 3F-H). Compound 527 did not show any inhibitory effects on blocking the activation of the receptors by their respective ligands (Figure 3E-H).

**Figure 3.**
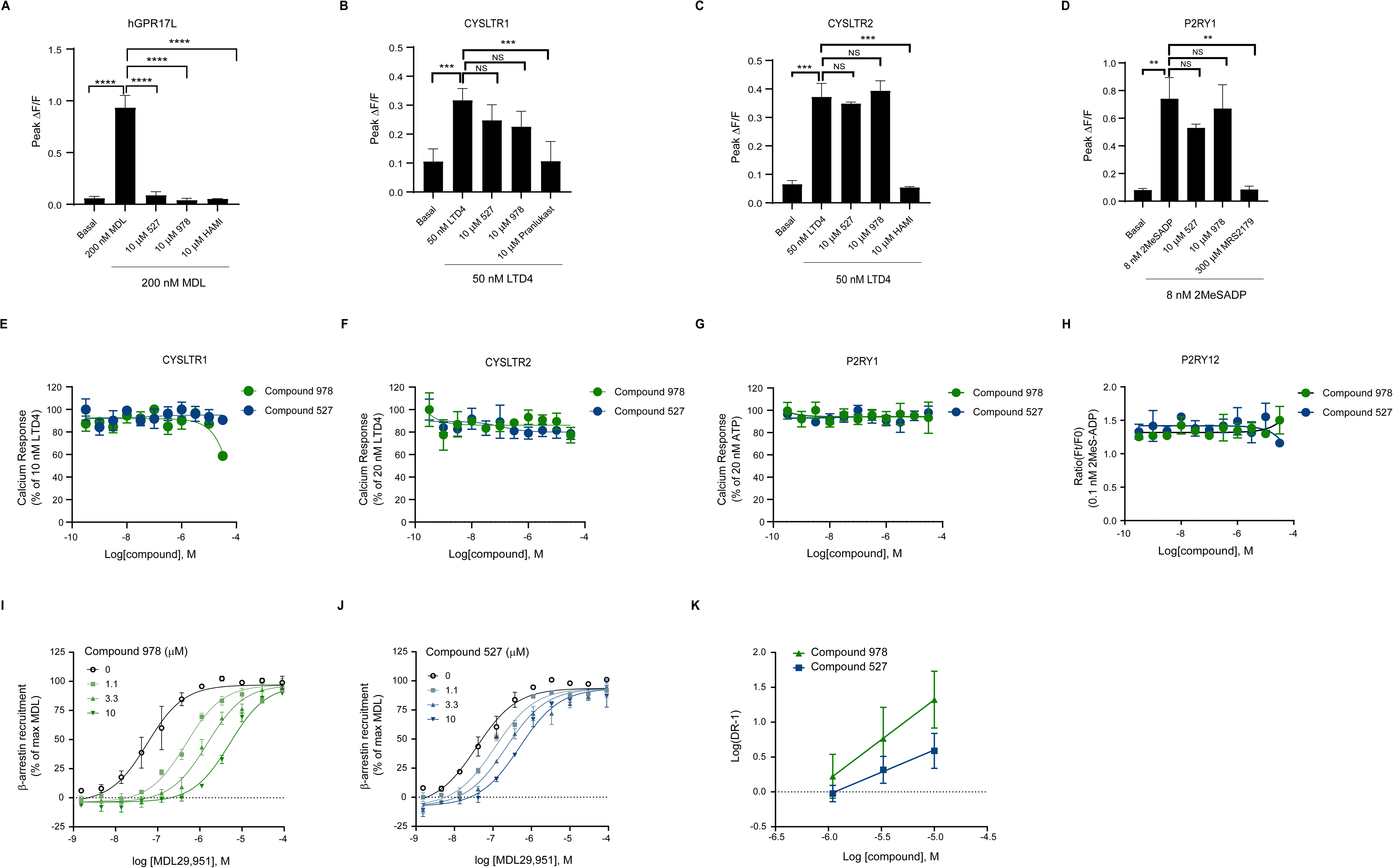
Selectivity of two hits against the receptors from phylogenetically close subfamilies of GPR17. Neuro-2a cell, a mouse neuroblastoma cell line, was transiently transfected with hGPR17L, hCysLTR1, hCysLTR2, and hP2RY1, and the antagonism of compounds against these receptors was assessed by calcium mobilization assay after stimulating with their respective agonist ligands. **A.** Neuro-2a cells expressing hGPR17 were pretreated with 10 µM of compounds 978, 527 and HAMI for 10 minutes before stimulated with 200 nM MDL. **B.** Neuro-2a cells expressing hCysLTR1 receptor were pretreated with 10 µM of compounds 978, 527 and pranlukast for 10 minutes before stimulated with 50 nM LTD4. **C.** Neuro-2a cells expressing hCysLTR2 receptor were pretreated with 10 µM of compounds 978, 527 and HAMI for 10 minutes before stimulated with 50 nM LTD4. **D.** Neuro-2a cells expressing hP2Y1 receptor were pretreated with 10 µM of compounds 978 and 527, and 300 µM MRS2179 for 10 minutes before stimulated with 8 nM 2MeSADP. Data (**A-D**) were analyzed by one-way ANOVA followed by Dunnett’s post hoc test where each test compound treatment condition was compared to vehicle treatment. * p < 0.05, ** p < 0.01, *** p < 0.001, **** p < 0.0001. Selectivity of two hits was also assessed by hCysLTR1, hCysLTR2, hP2RY1, hP2RY12 HEK293 stable cells. **E.** hCysLTR1 stable cell line was pretreated with the compounds for 10 min and then stimulated with 10nM LTD4. **F.** hCysLTR2 stable cell line was pretreated with the compounds for 10 min and then stimulated with 10nM LTD4. **G.** hP2RY1 stable cells were pretreated with the compounds for 10 min and then stimulated with 20nM ATP. **H.** hP2RY12 stable cells were treated with the compounds for 25 min and then stimulated with 0.1 nM 2MeS-ADP. The signals were detected by FlexStation. Schild regression analysis of two antagonists in β-arrestin recruitment assay. **I**. Dose response curve of MDL in the presence of increased concentration of compound 978 in β-arrestin recruitment assay. **J**. Dose response curve of MDL in the presence of increased concentration of compound 527 in β-arrestin recruitment assay. **K**. Schild plot of compounds 978 and 527 in β-arrestin recruitment assay. The slopes of compounds 978 and 527 were 1.15 and 0.64 respectively. The pA2 value of compounds 978 and 527 were 6.2 and 5.9 respectively.

The validation and characterization of compounds 978 and 527 suggested specific and selective antagonism of GPR17, but these assays do not provide information regarding the molecular mechanisms of antagonism. Therefore, the mode of action for each compound was investigated in β-arrestin recruitment assays by evaluating the effects of varying concentrations of test compounds on dose-response curves of MDL in 1321N1-GPR17 cells. To test the effects of compounds on GPR17-mediated β-arrestin recruitment, compounds 978 and 527 were added 10 minutes prior to addition of the agonist MDL in 1321N1-hGPR17 cells. β-arrestin recruitment was measured using the Cisbio HTRF β-arrestin recruitment assay. 1 µM, 3.3 µM, and 10 µM concentrations of compounds 978 (Figure 3I) and 527 (Figure 3J) caused concentration-dependent rightward shifts of the MDL concentration-response curve for GPR17-mediated β-arrestin recruitment. Schild regression analysis for compounds 978 and 527 yielded slopes of 1.15 and 0.64, respectively (Figure 3K). They were not significantly different than one, as demonstrated in 95% confidence intervals (Table 1). This suggests a mechanism of surmountable antagonism that is consistent with a competitive interaction between these compounds and MDL at the binding site of the receptor.

**Table 1.** Schild Analysis of compounds in calcium mobilization assay and β-arrestin recruitment assay.

| Compound | β-arrestin Schild Slope (95% CI) |
| --- | --- |
| 527 | 0.81 (0.62 – 1.02) |
| 978 | 1.13 (0.96 – 1.30) |

### Structure activity relationship of antagonists

After identification of compounds 978 and 527 as selective antagonists of GPR17, we engaged in exploratory structure-activity relationship (SAR) campaigns around each of the chemotypes. To that end, we developed synthetic routes that were amenable to diversification in most regions of the chemical structures. For the chemical series represented by 978, we started the synthesis with substituted phenyl hydrazines such as 1-chlorophenyl hydrazine (Figure 4A, a). These were condensed with cyclohexane-1,2-dione (Figure 4A, b) to provide a tetrahydrocarbozol-1-one (Figure 4A, c). whose ketone was then reduced using ammonium acetate and sodium cyanoborohydride to the corresponding aliphatic amine (Figure 4A, d). This amine served as a handle for the HATU-mediated coupling with diverse carboxylic acids (Figure 4A, e) to provide a range of structurally diverse tetrahydrocarbazolyl amides (Figure 4A, f). A modular synthesis was also developed for the 527 chemotype. The synthetic sequence (Figure 4B) started with the three-component coupling of thiophene-2-sulfonamide (Figure 4B, g), oxoacetic acid (Figure 4B, h), and phenyl boronic acid (Figure 4B, i) to provide central intermediate (Figure 4B, j). This has a carboxylic acid handle for HATU-mediated amide coupling with diverse anilines (Figure 4B, k).

**Figure 4.**
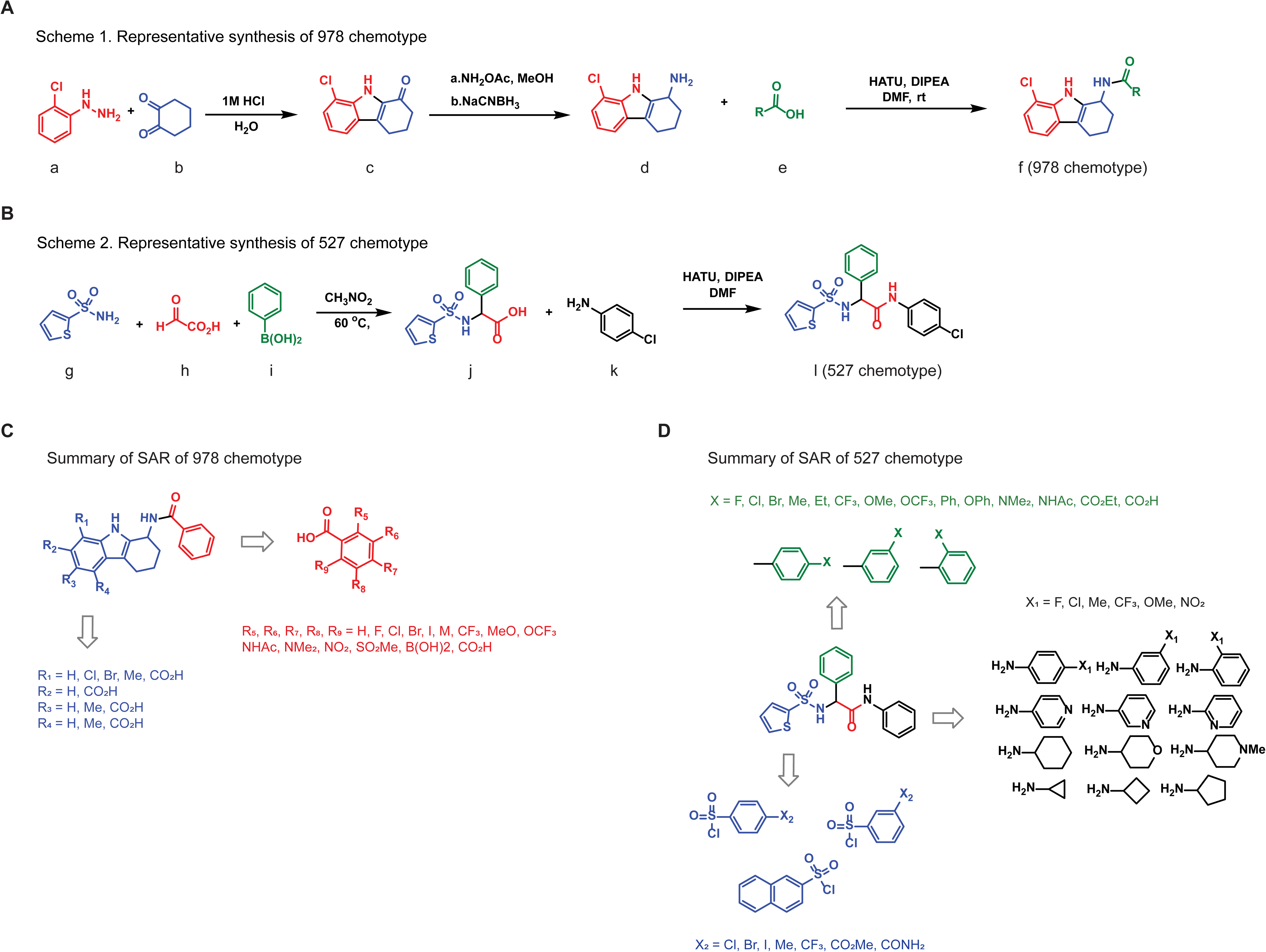
Schemes 1 and 2 are representative synthetic schemes to make analogs of chemotypes 978 and 572. Tables 1-2 show SAR around chemotype represented by 978. Tables 3-5 show SAR around chemotype represented by 572. IC_50_s are reported when the % inhibition is greater than 75%. The efficacy is the percentage inhibition at the highest compound concentration in the assay.

**Table 2.**
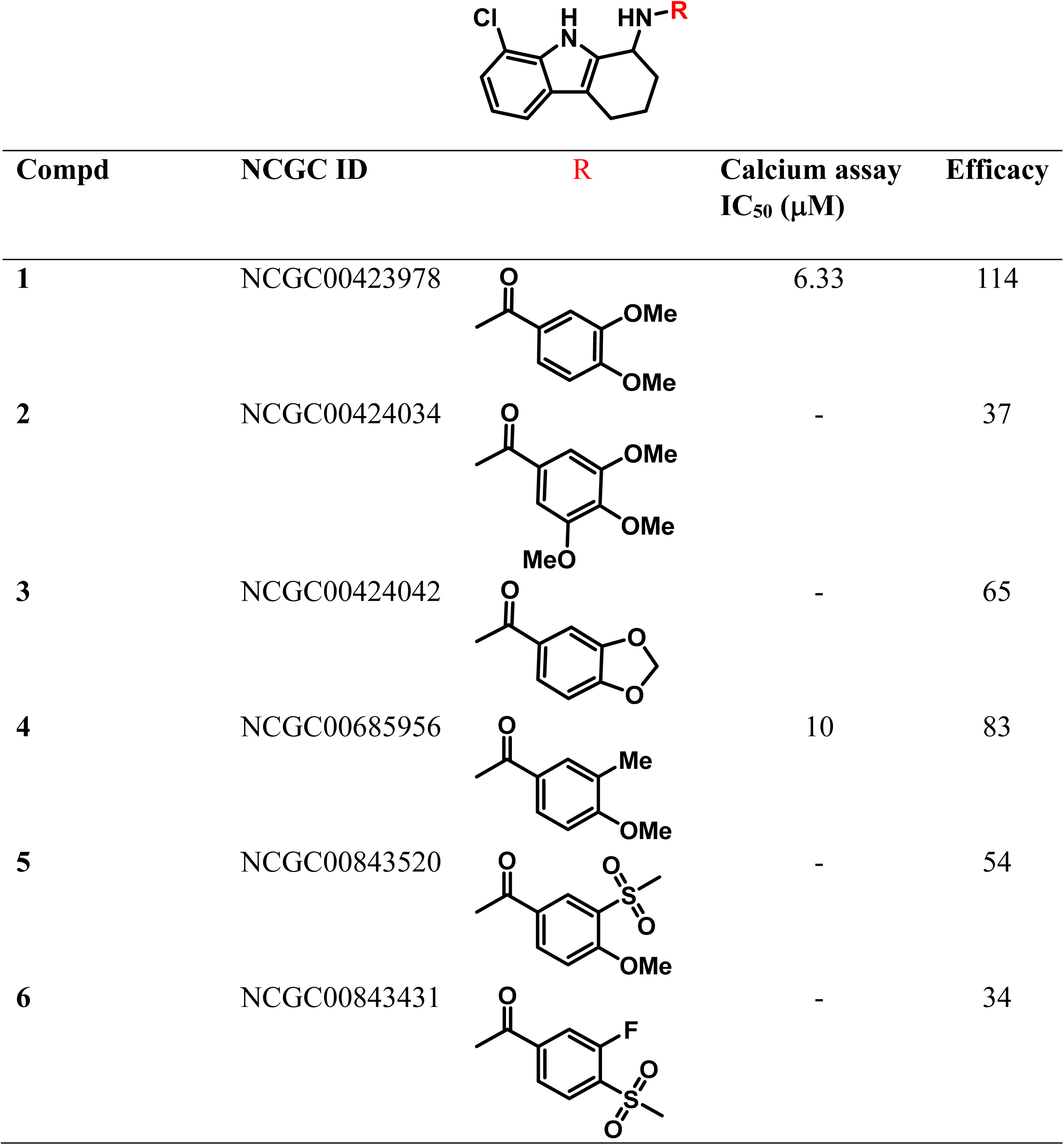
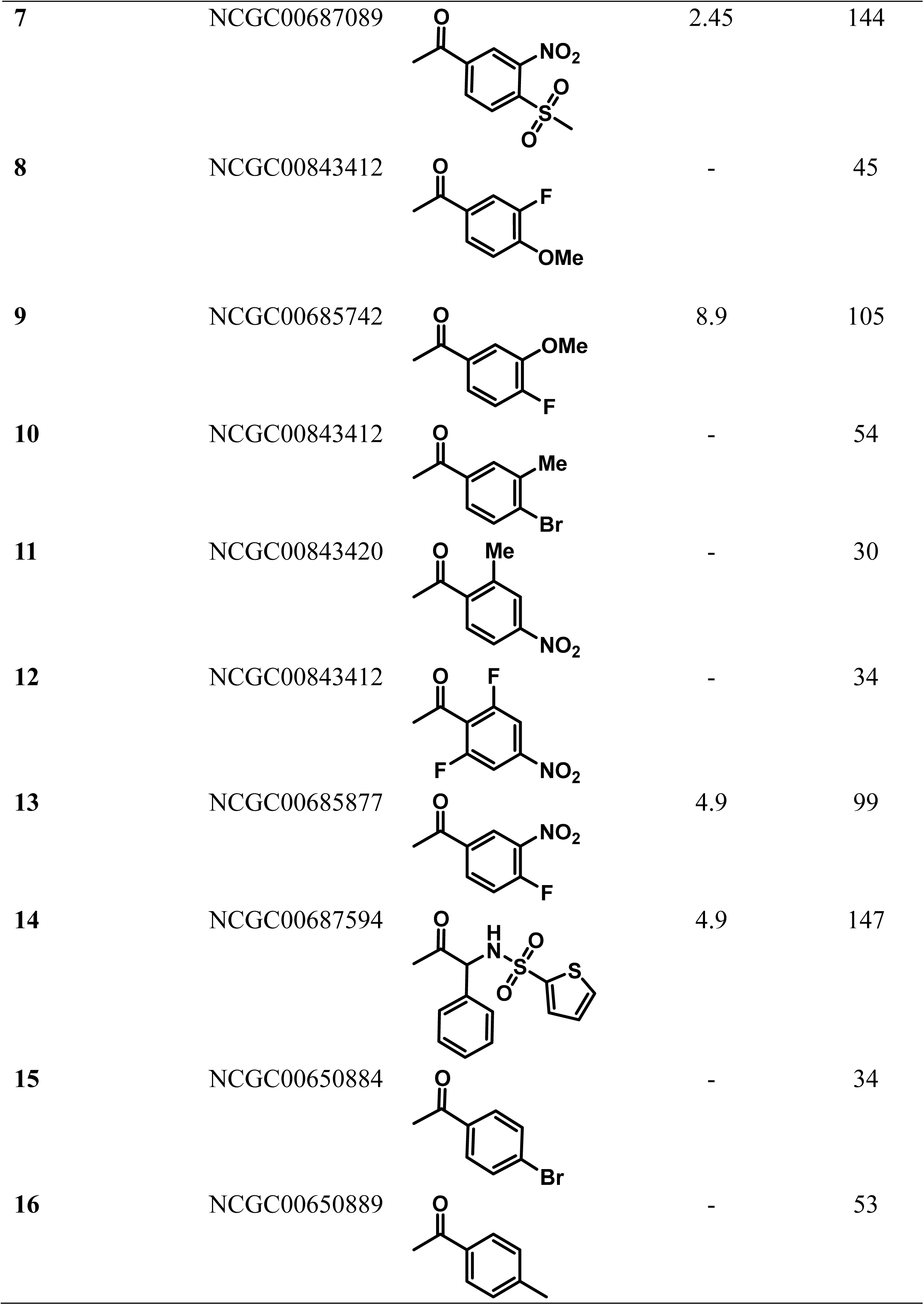
978 chemotype amide SAR.

| Compd | NCGC ID | R | Calcium assay<br>IC<sub>50</sub> (μM) | Efficacy |
| --- | --- | --- | --- | --- |
| 1 | NCGC00423978 |  | 6.33 | 114 |
| 2 | NCGC00424034 |  | - | 37 |
| 3 | NCGC00424042 |  | - | 65 |
| 4 | NCGC00685956 |  | 10 | 83 |
| 5 | NCGC00843520 |  | - | 54 |
| 6 | NCGC00843431 |  | - | 34 |
| 7 | NCGC00687089 |  | 2.45 | 144 |
| 8 | NCGC00843412 |  | - | 45 |
| 9 | NCGC00685742 |  | 8.9 | 105 |
| 10 | NCGC00843412 |  | - | 54 |
| 11 | NCGC00843420 |  | - | 30 |
| 12 | NCGC00843412 |  | - | 34 |
| 13 | NCGC00685877 |  | 4.9 | 99 |
| 14 | NCGC00687594 |  | 4.9 | 147 |
| 15 | NCGC00650884 |  | - | 34 |
| 16 | NCGC00650889 |  | - | 53 |

**Table 3.** 978 chemotype tetrahydrocarbazol SAR.

| Compd | NCGC ID | R<sub>1</sub> | R<sub>2</sub> | R<sub>3</sub> | R<sub>4</sub> | R<sub>5</sub> | R<sub>6</sub> | R<sub>7</sub> | Calcium assay<br>IC<sub>50</sub> (μM) | Efficacy |
| --- | --- | --- | --- | --- | --- | --- | --- | --- | --- | --- |
| 17 | NCGC00423976 | H | H | H | H | OMe | OMe | H | 6.64 | 81 |
| 18 | NCGC00841699 | Br | H | H | H | OMe | OMe | H | 3.99 | 120 |
| 19 | NCGC00843841 | Br | H | H | H | OMe | H | OMe | - | 57 |
| 20 | NCGC00843850 | Me | H | Me | H | OMe | OMe | H | 11.2 | 86 |
| 21 | NCGC00843432 | Me | H | Me | H | OMe | H | OMe | - | 71 |
| 22 | NCGC00846053 | Me | H | Me | Me | OMe | H | OMe | 12.6 | 84 |
| 23 | NCGC00842971 | CO<sub>2</sub>H | H | H | H | OMe | OMe | H | inactive |  |
| 24 | NCGC00841530 | H | CO<sub>2</sub>H | H | H | OMe | OMe | H | inactive |  |
| 25 | NCGC00842489 | H | H | CO<sub>2</sub>H | H | OMe | OMe | H | inactive |  |
| 26 | NCGC00842955 | Cl | H | CO<sub>2</sub>H | H | OMe | OMe | H | inactive |  |
| 27 | NCGC00842648 | Cl | H | H | CO<sub>2</sub>H | OMe | OMe | H | inactive |  |

We explored the SAR around chemotype 978 (Figure 4C). Representative synthesized analogs around the 978 chemotype are presented in Tables 2 and 3. Their activities are reported in the calcium assay. The IC_50_ is reported only when the activity at the highest concentration is greater than 75%. The efficacy is reported as a percentage; it is essentially the curve-fitted activity at the highest concentration of the assay. Table 2 shows the SAR at the amide region where a variety of electron donating groups (entries 2, 3, 4, 16) as well as withdrawing substitutions (entries 6, 7, 11, 12, 13, 15) are explored. Analogs with a combination of electron withdrawing and donating groups are also shown in entries 5, 6, 8, 9, 10, 11. An analogue that borrows features from the chemotype represented by 527 is shown in entry 14. Only 6 analogs, including the hit 978, in Table 2 show a maximum response greater than 75% inhibition and had an IC_50_ ≤ 10 μM. Table 3 shows the SAR in the aromatic ring present within the tetrahydro carbazole core.

The R_1_ substituent which was chloride in hit 978 was replaced with hydrogen (17), bromide (entries 18, 19) and a methyl group (entries 20-22). In this group of compounds, the bromide 18 with an IC_50_ ∼ 4 μM had an activity that was slightly better than hit 978. A carboxylic acid group was also systematically walked around the aromatic (R_1_ to R_4_) ring to mimic the carboxylic acids present in MDL with the hope of interacting with polar positively charged residues in GPR17. However, these compounds were inactive in the calcium assay.

We also explored the SAR around chemo type 527 (Figure 4D). Representative analogs are presented in Tables 4-6. Table 4 showcases an exploration of substitution at the central phenyl ring. Entries 29-31 explore ortho substitutions, 32-34 meta substitutions, and 35-44 para substitutions. Although a variety of electron withdrawing (fluoro, chloro, ester, acid, amide), electron donating (methyl, methoxy, alkyl), and lipophilic groups were explored, none of them showed inhibitory activity that were discernibly more important than the starting compound 527. Tables 5 and 6 show representative compounds that explored SAR around the western sulfonamide aromatic ring and eastern amide. Unfortunately, the structural diversity explored in these regions did not lead to an analog with better activity than compound 527. Thus, our medicinal chemistry efforts suggested that both chemotypes might be associated with SARs that are relatively flat.

**Table 4.**
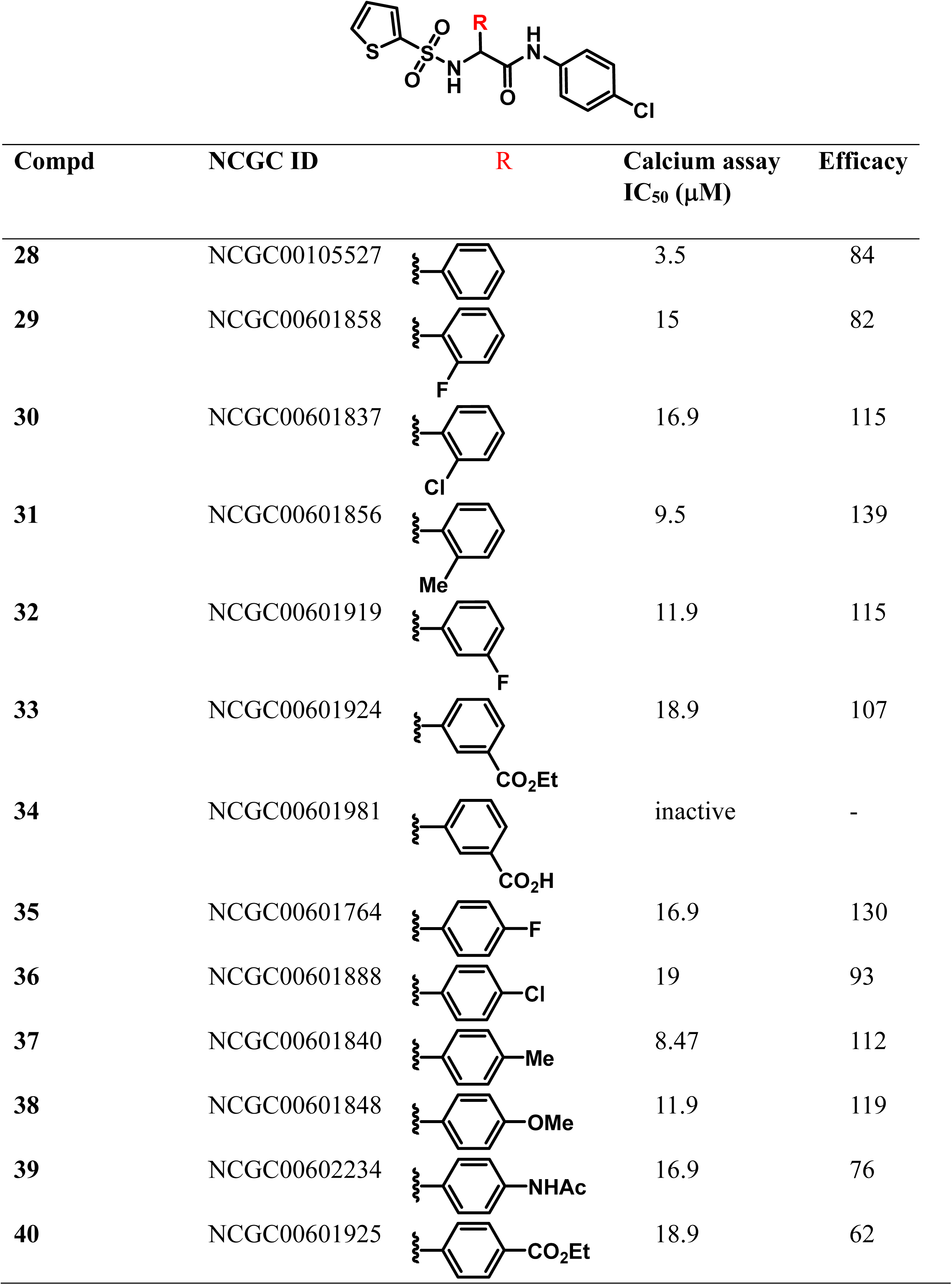
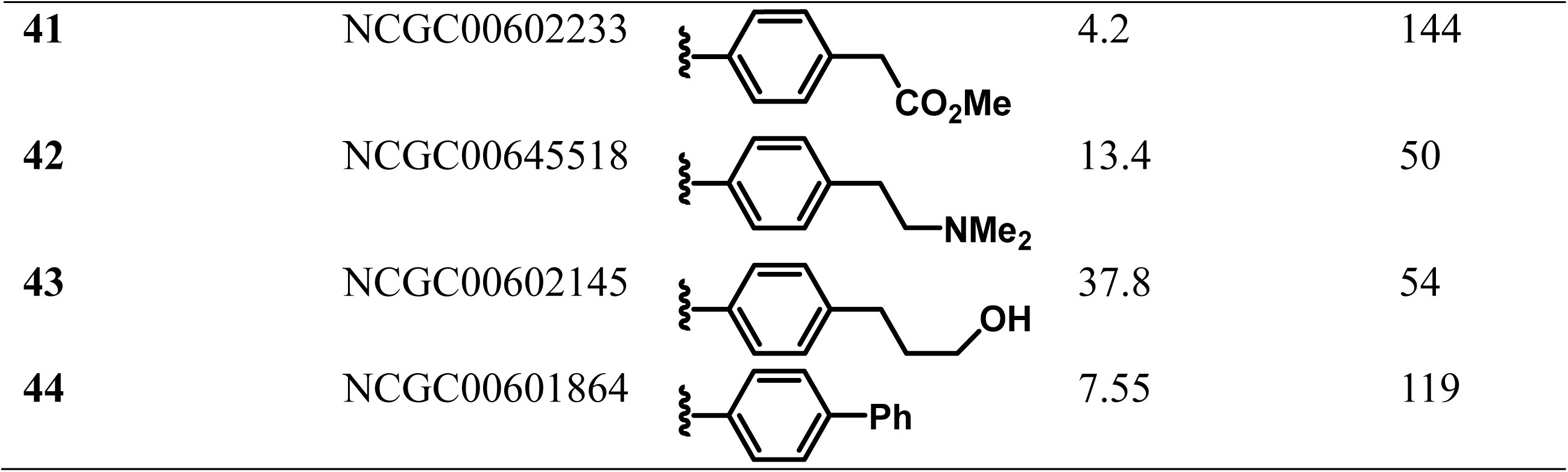
527 Chemotype central ring SAR.

| Compd | NCGC ID | R | Calcium assay<br>IC<sub>50</sub> (μM) | Efficacy |
| --- | --- | --- | --- | --- |
| 28 | NCGC00105527 |  | 3.5 | 84 |
| 29 | NCGC00601858 |  | 15 | 82 |
| 30 | NCGC00601837 |  | 16.9 | 115 |
| 31 | NCGC00601856 |  | 9.5 | 139 |
| 32 | NCGC00601919 |  | 11.9 | 115 |
| 33 | NCGC00601924 |  | 18.9 | 107 |
| 34 | NCGC00601981 |  | inactive | - |
| 35 | NCGC00601764 |  | 16.9 | 130 |
| 36 | NCGC00601888 |  | 19 | 93 |
| 37 | NCGC00601840 |  | 8.47 | 112 |
| 38 | NCGC00601848 |  | 11.9 | 119 |
| 39 | NCGC00602234 |  | 16.9 | 76 |
| 40 | NCGC00601925 |  | 18.9 | 62 |
| 41 | NCGC00602233 |  | 4.2 | 144 |
| 42 | NCGC00645518 |  | 13.4 | 50 |
| 43 | NCGC00602145 |  | 37.8 | 54 |
| 44 | NCGC00601864 |  | 7.55 | 119 |

**Table 5.** 527 chemotype sulfonamide SAR.

| Compd | NCGC ID | R | Calcium assay<br>IC<sub>50</sub> (μM) | Efficacy |
| --- | --- | --- | --- | --- |
| 45 | NCGC00650036 |  | 18.9 | 90 |
| 46 | NCGC00650047 |  | 26.7 | 138 |
| 47 | NCGC00650052 |  | 21.2 | 91 |
| 48 | NCGC00650046 |  | inactive | - |
| 49 | NCGC00650049 |  | 16.9 | 120 |
| 50 | NCGC00650043 |  | - | 58 |
| 51 | NCGC00650039 |  | inactive | - |
| 52 | NCGC00650038 |  | inactive | - |

**Table 6.**
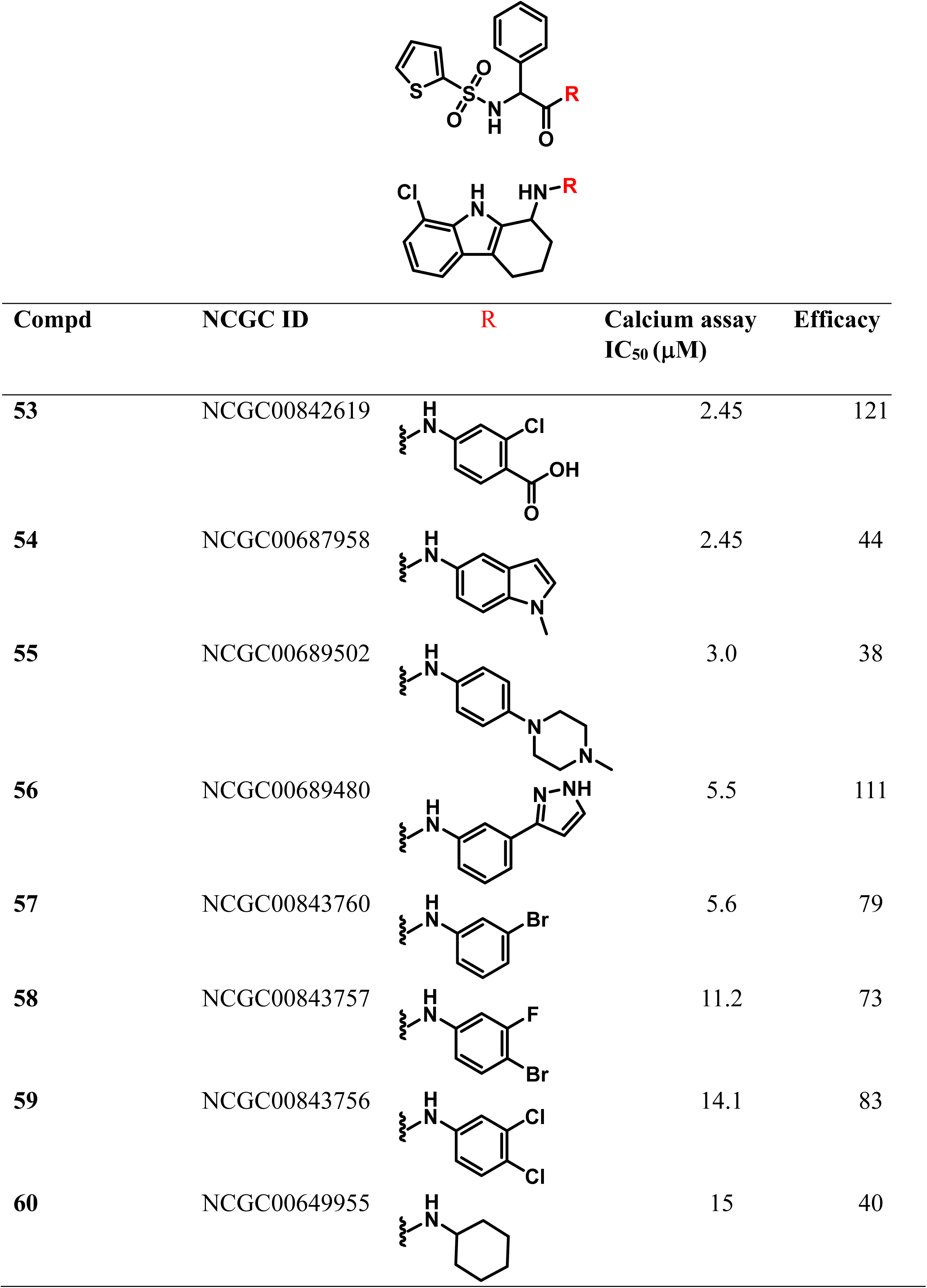
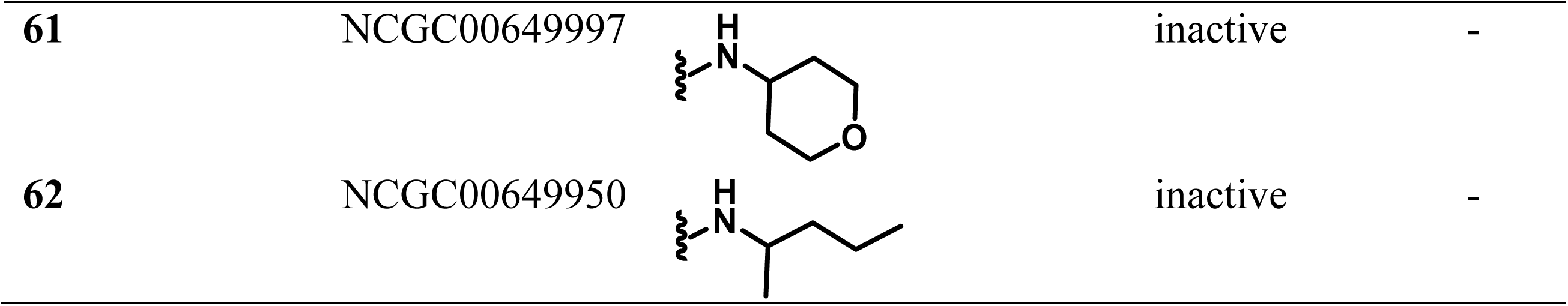
527 chemotype amide SAR.

### Homologous model of antagonists with GPR17

To simulate the interaction of compounds 978 and 527 with GPR17, a predicted structure of human GPR17 was downloaded from AlphaFold Protein Structure Database (https://alphafold.ebi.ac.uk/). The seven-transmembrane structure of GPR17 formed a ligand binding site on the extracellular side, which was basic (Fig.5A and 5B). Molecular docking was carried out using the Dock module in the Molecular Operation Environment (MOE) by Chemical Computing Group (www.chemcomp.com). The protein structure was allowed a certain degree of flexibility by the Induced Fit algorithm. The predicted pose of the known agonist MDL formed a salt bridge with ARG^255^, and a hydrogen bond with GLU^187^ (Fig. 5C). The phenyl ring and thiophenyl ring in GPR17 inhibitor compound 527 were predicted to be stacked in the binding site, whereas sulphone and amide oxygen atoms were involved in hydrogen bonding interactions with LYS^188^ and ARG^255^ (Fig. 5D). The compound 978 interacted with GPR17 largely through hydrophobic interactions, by burying its tricyclic group into the narrow binding pocket (Fig. 5E).

**Figure 5.**
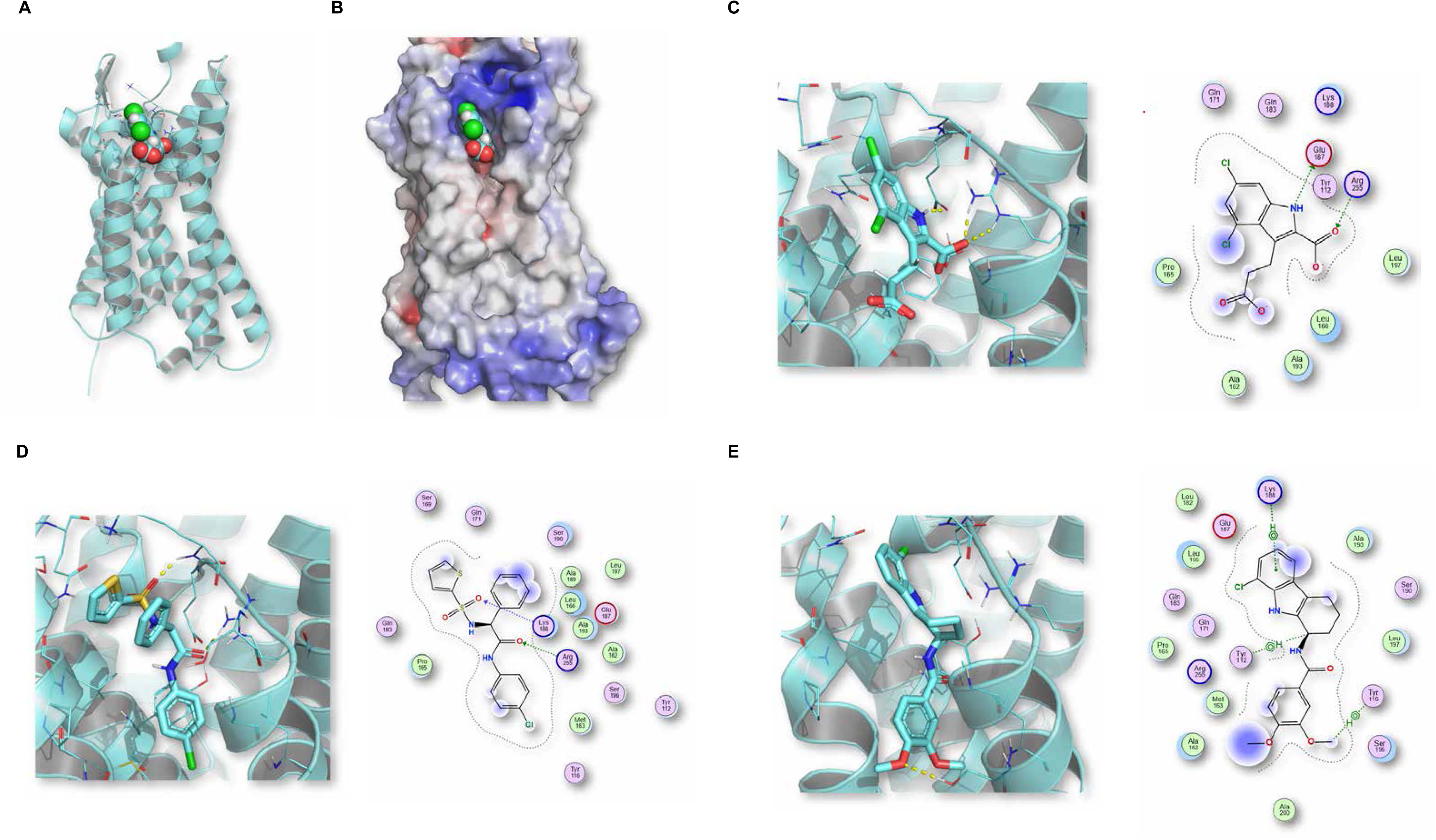
Homogeneous model of GPR17 docking with MDL and two antagonists. The predicted structure of human GPR17 with the agonist MDL docked in the ligand binding site: (A) Cartoon presentation (B) Electrostatic potential surface (blue color indicates electron-deficient surface, and red surface is electron rich) of the seven-transmembrane structure. The predicted ligand binding poses by molecular docking for (C) MDL, (D) compound 527 and (E) compound 978. The binding modes were displayed in both 3D structural presentation (Left panel) and 2D molecular interaction maps (Right panel).

### GPR17 antagonist effects on cAMP signaling and regulation of GLP-1 secretion in GLUTag cells

GPR17 is expressed in enteroendocrine cells of the gastrointestinal tract and regulates nutrient-stimulated GLP-1 secretion and thereby modulates glucose-stimulated insulin secretion and glucose homeostasis ^12^. Such effects are thought to be mediated by GPR17 Gi/o coupling ^12^. The GLUTag cell line is a GLP-1 secreting enteroendocrine cell model that has been extensively used to investigate GPCR regulation of GLP-1 secretion ^33–35^. We evaluated compounds 978 and 527, as well as some analogues from SAR studies (Supplemental figure 3), for effects on GPR17-mediated cAMP signaling and GLP-1 secretion in GLUTag cells.

The putative GPR17 antagonists were first tested for effects on GPR17-mediated cAMP signaling modulation. Forskolin stimulation of hGPR17L expressing GLUTag cells elevated cAMP levels from 0.59 ± 0.04 nM to 4.7 ± 0.36 nM. Forskolin-stimulated cAMP production was inhibited by the activation of GPR17 with MDL to 3.5 ± 0.18 nM, representing a 29 ± 3% inhibition of forskolin stimulated cAMP, consistent with Gi/o coupling of GPR17 in GLUTag cells. The compounds (10 µM) were evaluated for the ability to attenuate MDL-mediated inhibition of forskolin-stimulated cAMP (Figure 6A). Compound 978 and its analogues (compounds 419, 850, and 793), compound 527, and the positive control HAMI significantly attenuated the 100 nM MDL inhibition of forskolin-stimulated cAMP in GLUTag cells expressing hGPR17L. Surprisingly, 527 analogues (compounds 518 and 480) had no effect on the MDL inhibition of forskolin-stimulated cAMP (Figure 6A). The attenuating effects on cAMP inhibition appear to be specific to GPR17-mediated inhibition of cAMP, as no attenuating effects were observed for any test compounds when forskolin-stimulated cAMP was inhibited by activating endogenous somatostatin receptors (Supplemental figure 4). Therefore, 978, 419, 850, 793, 527, and HAMI demonstrated antagonist activity with respect to GPR17-mediated cAMP signaling modulation in GLUTag cells.

**Figure 6.**
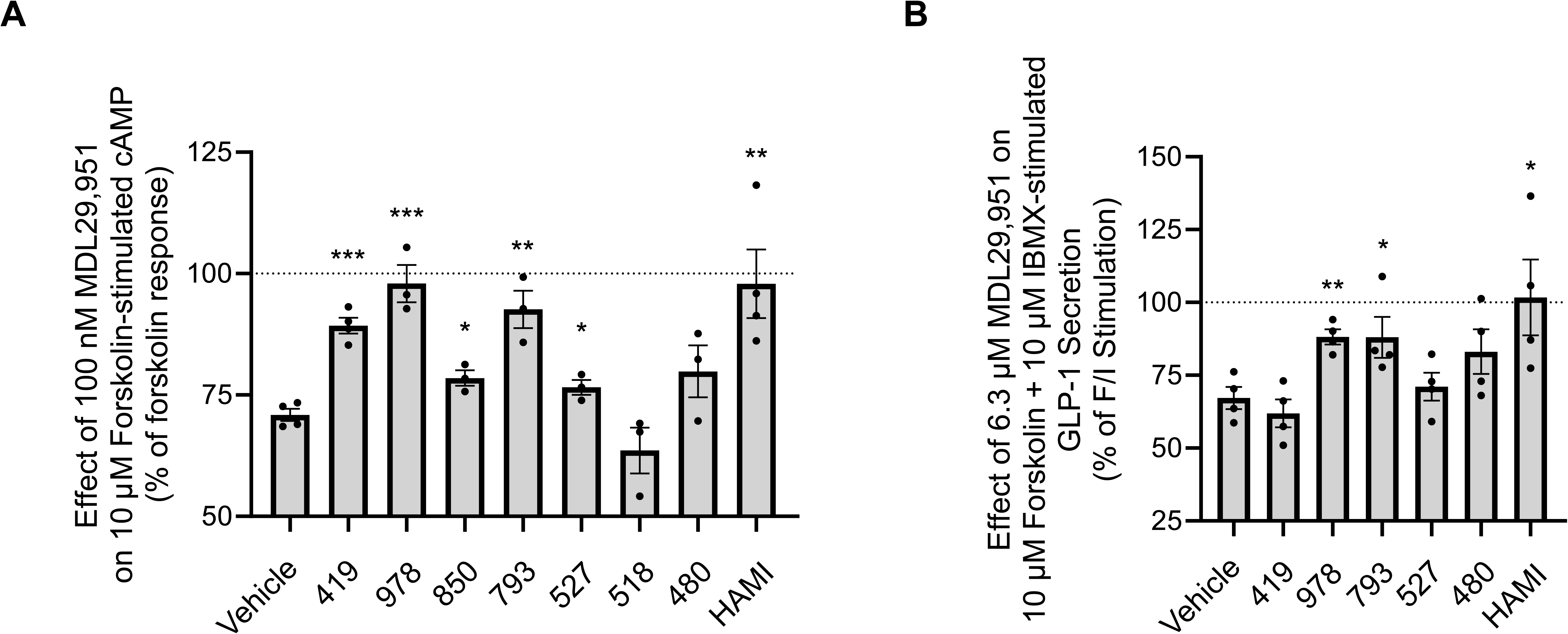
Confirmation of GPR17 antagonist activity for cAMP signaling and GLP-1 secretion modulation in GLUTag cells. **A.** Evaluation of test compounds for effects on 100 nM MDL-mediated inhibition of 10 µM forskolin-stimulated cAMP in GLUTag cells expressing hGPR17L. Data were normalized to the forskolin response within each test compound treatment condition and represent mean ± SEM of three to four independent experiments performed in triplicate wells. Data were analyzed by unpaired t-test where each test compound treatment condition was compared to vehicle treatment. * p < 0.05, ** p < 0.01, *** p < 0.001. **B.** GLP-1 secretion assay in GLUTag cells expressing hGPR17L. Test compounds were tested for effects on 6.3 µM MDL-mediated inhibition of 10 µM forskolin and 10 µM IBMX-stimulated GLP-1 secretion in GLUTag cells expressing hGPR17L. Data were normalized to the forskolin and IBMX response within each test compound treatment condition and represent mean ± SEM of four independent experiments performed in triplicate wells. Data were analyzed by unpaired t-test where each test compound treatment condition was compared to vehicle treatment. * p < 0.05, ** p < 0.01.

The compounds were also evaluated for effects on GPR17-mediated regulation of GLP-1 secretion. GLUTag cells that were transduced with adenovirus for expression of hGPR17L had 19.7 ± 2.6 pM GLP-1 in the supernatant under basal conditions. Stimulating hGPR17L expressing GLUTag cells with 10 µM forskolin and 10 µM IBMX caused over 3-fold increase of GLP-1 secretion (65.3 ± 8.0 pM vs 19.7 ± 2.6 pM at baseline) (Supplemental figure 5). Activation of hGPR17L with 6.3 µM MDL inhibited the forskolin and IBMX-stimulated GLP-1 secretion to 50.1 ± 4.6 pM (i.e., 33 ± 8% reduction), consistent with GPR17-mediated negative regulation of GLP-1 secretion (Supplemental figure 5). The compounds were evaluated for the ability to attenuate the MDL-mediated inhibition of forskolin and IBMX-stimulated GLP-1 secretion (Figure 6B). Compound 978, 793, and HAMI significantly attenuated MDL-mediated inhibition of GLP-1 secretion in GLUTag cells expressing hGPR17L. In summary, 10 µM compound 978 and its analogue compound 793 significantly attenuated MDL-mediated inhibition of cAMP and GLP-1 secretion in GLUTag cells expressing hGPR17L. These effects are consistent with GPR17 antagonist activity and suggest potential for targeting GPR17 with small molecule antagonists to enhance GLP-1 secretion.

## Discussion

In our efforts to explore new chemical space for GPR17 antagonists, we screened over 300K compounds. Two antagonists with novel scaffolds (compound 978 and 527) were successfully identified and validated in multiple assays. The compounds from both scaffolds effectively inhibited multiple downstream signaling pathways of GPR17 with high selectivity over other closely related receptors. Our findings provide new insights into the molecular mechanisms of GPR17 antagonism and offer valuable pharmacological tools for advancing therapeutic strategies in diabetes and obesity.

GPR17 is an orphan GPCR, and its putative endogenous ligand has not yet been conclusively validated. GPR17 is in an intermediate position between the P2RY and CysLTR families (27). Initial reports indicated that endogenous ligands from both families (UDP, UDP-glucose, LTC4 and LTD4) activated GPR17 in both cAMP accumulation assay and calcium mobilization assay ^16^. However, subsequent studies by other research groups failed to confirm these findings ^17,22,36^. Harrington et al. ^22^ reported that oxysterols, particularly 24(S)-hydroxycholesterol (24S-HC), could be the physiological activators of GPR17. Notably, 24S-HC, the major brain cholesterol metabolite, is an agonist of liver X receptors ^37^ and a positive allosteric modulator of NMDA receptor ^38^. Analysis of the data presented in their study suggests 24S-HC behaves more like an allosteric modulator instead of an endogenous agonist ^22^.

However, we were not able to confirm that 24S-HC either activated GPR17 or potentiated the activity of MDL in both cAMP accumulation assay and calcium mobilization assay (supplemental figure 6). To identify the potential lipid ligands of receptor, we cherrypicked and screened a collection of 745 lipid-like compounds from our in-house library. However, we can’t identify any compounds activating the receptor (data now shown). But we can’t exclude the possibility that the lipid-like molecules are the endogenous ligands of receptor,

It is intriguing that while endogenous ligands from closely related P2RY and CYSLTR families were unable to activate GPR17, antagonists from the CYSLTR subfamily can completely (pranlukast, HAMI and BayCysLT2) or partially (monelukast) block the MDL-stimulated receptor activation of GPR17 ^21^. In contrast, antagonists from the P2RYs family (ticagrelor and cangrelor) failed to block MDL-stimulated receptor activation ^22,25^. This phenomenon was also observed in our studies (data not shown). In the calcium mobilization assay, we found that pranlukast, HAMI, BayCysLT2 and zafirlukast showed full antagonist activity, montelukast and tomelukast showed partial antagonist activity, while MK571 showed no antagonist activity. The CYSLTR2 selective antagonists (HAMI and BayCysLT2) showed better antagonist activity than CYSLTR1 selective antagonists (montelukast and MK571) at GPR17.

Like previous studies, there is no antagonist activity in P2RY12 antagonists (clopidogrel, prasugrel and ticlopidine). The cross activity of these antagonists indicates that GPR17 might be more closely related to the CYSLTR family than the P2RY family. Besides these promiscuous and weak antagonists, more potentGPR17 antagonists (I-116 ^26^, UCB6651 ^27^, and PSB-22269 ^28^) were developed by Müller’s group. I-116 and UCB6651 were derived from the same scaffold.

UCB6651 showed nanomolar range potency in blocking GPR17 activation, while it is inert in four P2Rys (P2RY1, P2RY12, P2YR13 and P2RY14) and one CYSLTR (CYSLTR1). However, the effects of UCB6651 on CYSLTR2, a closely related receptor were not reported ^27^. PSB-22269, an anthranilic acid derivative, showed nanomolar range potency in the binding assay and the functional assays, while selectivity data were not reported ^28^. Compound 978 and 527 we discovered showed high selectivity for GPR17, and no activities was observed in P2RYs (P2RY1 and P2RY12) and CYSLTRs (CYSLTR1 and CYSLTR2). Two novel selective scaffolds we identified will serve as promising starting points for further optimization, potentially advancing the development of targeted therapies with improved efficacy and safety for treating diabetes and obesity.

Although the identity of the endogenous ligand of GPR17 is still controversial and elusive, the synthetic small molecule agonist MDL was consistently validated by multiple groups including us ^12,13,22^. After activation by MDL, GPR17 was able to couple to major G protein and β-arrestin signaling pathways ^21^. As downstream signaling pathways were equally activated, MDL was considered as an unbiased agonist for the receptor. In our recent study, we demonstrated functional selectivity or biased signaling of GPR17 variants identified from metabolic disease patients ^13^. Eight nonsynonymous *GPR17* variants showed diverse downstream signaling profiles after being activated by MDL. Compared to the wildtype receptor, the GPR17-V96M variant showed biased Gαi/o signaling, and the GPR17-D105N variant showed biased Gαi/o and β-arrestin signaling pathways. GPCR biased signaling has important implications for drug development, as targeting biased signaling can result in better therapeutic effects with fewer adverse effects ^39^. The diverse downstream signaling profiles of GPR17 variants in metabolic disorders might indicate the importance of biased signaling in the disease and possibility of development of biased ligands to probe the different downstream signaling of receptor.

In conclusion, we identified two novel and selective antagonists (compounds 978 and 527) of GPR17 through high-throughput screening. Both antagonists blocked GPR17 downstream Gαi/o, Gαq and β-arrestin signaling with high selectivity for GPR17, but not the closely related purinergic and cysteinyl leukotriene receptors. One antagonist (978) and it analog (793) attenuated GPR17 signaling and promoted gut hormone, glucagon-like peptide-1 (GLP-1), secretion in enteroendocrine cells. Our findings suggested that antagonizing GPR17 might be a promising therapy for metabolic disorders.

## Materials and Methods

### Chemicals, libraries, and cell lines

The following four compounds were purchased in Tocris Bioscience (Minneapolis, MN): MDL29951 (Cat# 0467), ATP (Cat# 3245), L-692,585 (Cat# 2261), PF 05190457 (Cat# 6350).

Leukotriene D4 (LTD4, Cat# 20310) and 2-Methylthioadenosine diphosphate (2MeS-ADP, Cat# 21230) were purchased from Cayman Chemical (Ann Arbor, Michigan). HAMI 3379 was purchased from MedChemExpress (Monmouth Junction, NJ). Recombinant Human FGF2 (Cat# 233-FB-010) was purchased from R&D Systems (Minneapolis, MN). FIIN-2 (Cat# S7714) was purchased from Selleck Chemicals (Houston, TX). Five in-house libraries were screened, including Sytravon (a retired Pharma screening collection that contains a diversity of novel small molecules, NCATS), Genesis (an in-house high-value and modern chemical library, NCATS), NPC (NCATS Pharmaceutical Collection, NCATS), Kinase inhibitor library (an in-house collection of kinase inhibtors, NCATS) and LOPAC (1,280 Pharmacologically Active Compounds Sigma-Aldrich, St. Louis, MO) libraries. PathHunter U2OS GPR17 β-arrestin cell line Catalog# (Catalog# 93-0633C3), PathHunter U2OS FGFR1 cell line (Catalog# 93-1090C3) and PathHunter U2OS GHSR1a β-arrestin cell line (Catalog#93-0242C3) were purchased from DiscoverX/Eurofin (Fremont, CA). 1321N1 GPR17 stable cell line (Catalog# Cr1525-3) was purchased from Multispan Inc (Hayward, CA). HEK293-hGPR17S cell line (Catalog# CB-80400-301), HEK293-hCysLTR1 cell line (Catalog# CB-80400-299), HEK293-hCysLTR2 cell line (Catalog# CB-80400-300), HEK293-hP2RY1 cell line (Catalog# CB-80400-302), and -hP2RY12 cells (Catalog# CB-80300-303) were purchased from Codex Biosolutions Inc (Gaithersburg, MD).

### PathHunter β-arrestin recruitment assay

PathHunter U2OS GPR17 β-arrestin cells were maintained in the growth medium consisting of MEM (Thermo, # 11095080), 10% FBS (Thermo, # 10082) 1xPen/Strep, 1xGlutamine, 500µg/mL Geneticin (Thermo, # 10131035), and 250µg/mL Hygromycin B (Thermo, # 10687010).

For the high-throughput screen, cells were harvested with StemPro Accutase Cell Dissociation Reagent (Thermo, # A1110501), and seeded in 1536-well white flat bottom plates (Greiner, #789173-F) at the density of 500 cells/ 3µL/ well in AssayComplete Cell Plating Reagent 5 (Eurofin/DiscoverX, # 93-0563R5A). After incubated at 37°C, 5% CO_2_ overnight, 20 nL/well of library compounds were added to the columns 5-48 of each plate with a pintool transfer (Kalypsis, San Diego, CA). As the positive control, 20 nL/well of 10mM HAMI 3379 (66.7 µM final concentration) was transferred to column 3 and 4 of each plate. The microplates were incubated at room temperature for 10 minutes, and then ∼EC_80_ concentration of agonist MDL29951 (final concentration 2.0 µM) was added into the microplate. After stimulation, cells were incubated 37°C, 5% CO_2_ for 90 minutes. Finally, 1.5µL/ well of PathHunter detection reagent (DiscoverX# 93-0001L) was added to microplates with BioRAPTR FRD dispenser (Beckman Coulter, Brea, CA). The detection reagent was prepared by mixing Galacton Star Substrate, Emerald II solution and PathHunter buffer (supplied by the assay kit) together at 1:5:19 proportion accordingly prior to dispensing. The microplates were incubated at room temperature for 1 hour and then the luminescent signal was detected on ViewLux plate reader (PerkinElmer, Waltham, MA).

For the experiment of Schild regression analysis, 1321N1-GPR17 cells were seeded in 96-well plates. Cells were serum starved for 2 h and treated with varying concentrations of test compound and vehicle for 10 min in Stimulation Buffer 4 provided with the Cisbio HTRF Beta-Arrestin assay kit at room temperature. Cells were then treated with varying concentrations of the GPR17 agonist MDL in stimulation buffer 4 for 30 min at room temperature. The Cisbio HTRF Beta-Arrestin recruitment kit was then used to quantify the β-arrestin recruitment as specified in the manufacturer’s protocol. The HTRF signal was read using a Synergy 4 plate reader. Schild regression analysis was calculated using 4-variable concentration-response curves.

### Calcium mobilization assay

1321N1 GPR17 stable cell line was maintained in the growth medium consisting of DMEM (Thermo, # 10566016), 10% FBS (Thermo, # 10082) 1xPen/Strep, 1xGlutamine, 500µg/mL puromycin (Thermo, # A1113802).

For the high-throughput screen, cells were harvested with StemPro Accutase Cell Dissociation Reagent (Thermo, # A1110501), and seeded in 1536-well μclear black microplates (Greiner, #782097) at the density of 1000 cells/ 3µL/ well in DMEM medium. After incubated at 37°C, 5% CO_2_ overnight, 3 µL Calbryte™ 520NW dye-loading solution (AAT Bioquest, # 36319) was dispensed into each well. The dye-loading solution was prepared by mixing of 1xHHBS buffer, 10X Pluronic® F127 Plus and Calbryte™ 520NW stock solution (supplied by the assay kit) accordingly the manual from the kit. The dye-loading plates were incubated at 37°C, 5% CO_2_ for 30 minutes, and then incubated at room temperature for another 15-30 minutes. 20 nL/well of library compounds were transferred to the columns 5-48 of each plate by using Echo 525 acoustic dispenser (Beckman Coulter, CA). The microplates were incubated at room temperature for 10 minutes and then loaded on the FLIPR Penta High-Throughput Cellular Screening System (Molecular Devices, San Jose, CA). The baseline fluorescence signal was measured for 10 seconds. MDL (∼ EC_80_ concentration) was added to stimulate the cells, and then fluorescence signals were measured for 3 minutes at the 1-second interval. The relative fluorescence unit (RFU) was calculated by maximum signal – minimum signal.

For the experiment of Schild regression analysis, 1321N1-GPR17 cells were seeded into 384-well plates at a density of 7,500 cells/well. The following day, media was replaced with assay buffer (HBSS, 20 mM HEPES) and cells were loaded with Calcium 6 dye for 2 h at 37°C and 5% CO2. The cells were allowed to equilibrate to room temperature for 15 minutes and cells were incubated in test compound or vehicle for 10 minutes. Subsequently, MDL was added and plates were read every 0.5 s for 3 minutes on a Hamamatsu FDSS/μCell.

### cAMP accumulation assay

hGPR17-1321N1 cells were seeded at 3,000 cells/well into white, opaque 384-well plates and grown overnight at 37°C. The next day, cells were treated with varying concentrations of compound 527 or compound 978 for 10min at room temperature. Cells were treated with 30nM MDLin stimulation buffer (3 µM forskolin/500uM IBMX) for 1 hour at room temperature. HRTF assays were performed using the homogenous time-resolved fluorescence (HTRF) HiRange cAMP detection kit (Cisbio, Bedford, MA) according to the manufacturer’s instructions. Plates were incubated at room temperature for 30 minutes and FRET signals (665 and 620 nm) were read using Biotek Synergy Neo 2 plate reader (Agilent, Santa Clara, CA). HTRF signal was calculated as the ratio of signal from the 665 nm (acceptor) and 620 nm (donor) channels.

### ACTone cAMP assay

ACTOne-hP2RY12 cells (Codex Biosolutions, #CB-80300-303) were plated on a 384-well black clear plate at the density of about 12,000 cells/well in 20 ul culture medium. After incubated at 37°C, 5% CO_2_ overnight, 20 ul of 1x MP dye solution (Codex BioSolutions Inc) was loaded into each well. The plate was incubated at room temperature in the dark for 2 hours. Compounds were serially diluted (5x of the final concentration) in 1x DPBS solution. 10 ul of the diluted compounds were added into each well and incubated at room temperature in the dark for 25 min. The baseline fluorescent intensity (F0) of each well was recorded on FlexStation Multi-Mode Microplate Reader (Molecular Devices, San Jose, CA). 12.5 ul of 5x stimulation solution (0.1 nM 2MeS-ADP, 125 µM Ro20-1724 and 1.5 µM Iso in 1x DPBS) was then added and incubated at room temperature in the dark for 15 min. The fluorescent intensity (Ft) of each well was recorded again on FlexStation. The ratios of Ft/F0 were used to plot the dose response curves.

### Cisbio β-arrestin HRTF Assay

hGPR17-1321N1 cells were seeded at 30,000 cells/well into white, opaque 96-well plates and grown overnight. The next day, cells were serum starved then treated with varying concentrations of the stated compounds, or vehicle control, for 10min at room temperature. Cells where then treated with varying concentrations of MDL29,951 for 30min at room temperature. Cisbio manufactures protocol was then followed for the remainder of the experiment. HTRF signal was calculated as the ratio of signal from the 665 nm (acceptor) and 620 nm (donor) channels.

### GLUTag cell GLP-1 secretion assay

GLUTag cells were maintained in Dulbecco’s Modified Eagle Medium (DMEM), supplemented with 10% fetal bovine serum, and 1% penicillin-streptomycin in a humidified incubator at 37°C and 5% CO2. For adenovirus transduction, 75,000 cells/well GLUTag cells were seeded into poly-D-lysine coated 96-well plates. The next day, cells were transduced with Ad-CMV-HA-hGPR17L-EF1α-mCherry or Ad-EF1α-mCherry control adenovirus (625-1250 viral particles per cell at time of seeding) as indicated. After approximately 30 h, adenovirus-containing media was changed to serum-free growth media for overnight serum starvation. The following day, GLUTag cells were used for cAMP signaling assays or GLP-1 secretion assays.

GLUTag cells were transduced with adenovirus as specified above and then tested for cAMP signaling. Media was removed and cells were washed with cAMP stimulation buffer (Hanks balanced salt solution (HBSS), 20 mM HEPES, pH 7.4) and then incubated in cAMP stimulation buffer for 30 min at 37°C, 5% CO2. Cells were pre-treated with antagonist for 15 min prior to 30 min treatment with agonist and forskolin in the presence of 500 µM 3-isobutyl-1-methylxanthine (IBMX) at 37°C and 5% CO2. After treatment, stimulation buffer was decanted, and cAMP lysis buffer (50 mM HEPES, 10 mM CaCl2, 0.35% Triton X-100, and 500 µM IBMX) was added and shaken at 700 rpm for 1 h. Lysate was transferred to a 384-well plate and the cAMP levels were quantified using the Cisbio HTRF cAMP (Gi kit) according to the manufacturer’s protocol.

GLUTag cells were transduced with adenovirus as specified above and then tested for GLP-1 secretion. Media was removed and cells were washed two times with GLP-1 secretion buffer (138 mM NaCl, 4.5 mM KCl, 4.2 mM NaHCO3, 2.5 mM CaCl2, 1.2 mM MgCl2, 10 mM HEPES, 1 mM glucose, and 0.1% bovine serum albumin, pH 7.4). Cells were stimulated for 2 h with indicated treatments diluted in GLP-1 secretion buffer for a total volume of 200 µl/well at 37°C and 5% CO2. Plates were then centrifuged at 300xg for 1 min at 4°C and 75 µl of the supernatant was removed and frozen at −80°C until subsequent GLP-1 quantification. Total GLP-1 was quantified using the Meso Scale Discovery (Rockville, MD) Total GLP-1 V-PLEX kit (K1503PD) according to manufacturer’s instructions.

### HTS Data Analysis

Analysis of activity of compounds was performed as previously described ^40^. Briefly, raw plate read for each compound was first normalized relative to the HAMI control wells and DMSO control wells as follows: % Activity = ((RLU_compound_ – RLU_DMSO_)/(RLU_DMSO_ – RLU_HAMI_)) × 100, where RLU_compound_ denotes the compound well values, RLU_HAMI_ denotes the median value of the HAMI control wells, and RLU_DMSO_ denotes the median values of the DMSO-only wells, and then corrected by applying a NCGC in-house pattern correction algorithm using compound-free control plates (i.e., DMSO-only plates) at the beginning and end of the compound plate stack ^41^. For dose-response curves in confirmatory assay, counter assays and orthogonal assays of each compound were fitted to a four-parameter Hill equation ^42^ yielding concentrations of half-maximal activity (IC_50_) and maximal response (efficacy) values. Compounds were designated as Class 1–4 according to the type of dose–response curve observed ^40,43^. Curve classes are heuristic measures of data confidence, classifying concentration–responses based on efficacy, the number of data points observed above background activity, and the quality of fit. Compounds with −1.1, −1.2, −2.1, −2.2, −3.0 were considered active. Class 4 compounds were considered inactive. Compounds with other curve classes were deemed inconclusive. Active compounds were cherrypicked for follow up experiments.

