## Supplemental methods and figures for "Discovery of Novel and Selective GPR17 Antagonists as Pharmacological Tools for Developing New Therapeutic Strategies in Diabetes and Obesity"

**General Methods for Chemistry:** All air- or moisture-sensitive reactions were performed under positive pressure of nitrogen with oven-dried glassware. Anhydrous solvents such as dichloromethane, *N,N*-dimethylformamide (DMF), acetonitrile, methanol and triethylamine were purchased from Sigma-Aldrich. Preparative purification was performed on a Waters semi-preparative HPLC system. The column used was a Phenomenex Luna C18 (5 micron, 30 x 75 mm) at a flow rate of 45 mL/min. Purification was performed using **Method Acidic Standard Gradient** (10:90 MeCN with 0.1% TFA/ deionized water with 0.1% TFA ramped to 100% deionized water with 0.1% TFA) and **Method Basic Standard Gradient** (10:90 MeCN with 0.1% $NH_4OH$/ deionized water with 0.1% $NH_4OH$ ramped to 100% deionized water with 0.1% $NH_4OH$) over 8 minutes unless otherwise noted. Fraction collection was triggered by UV detection (220 nM). Analytical analysis was performed on an Agilent LC/MS (Agilent Technologies, Santa Clara, CA). Purity analysis was determined using a 7 minute gradient of 4% to 100% acetonitrile (containing 0.025% trifluoroacetic acid) in water (containing 0.05% trifluoroacetic acid) with an 8 minute run time at a flow rate of 1 mL/min. A Phenomenex Luna C18 column (3 micron, 3 x 75 mm) was used at a temperature of 50 °C using an Agilent Diode Array Detector.

Mass determination was performed using an Agilent 6130 mass spectrometer with electrospray ionization in the positive mode. $^1$H NMR spectra were recorded on Varian 400 MHz spectrometers. Chemical shifts are reported in ppm with non-deuterated solvent (DMSO-*d*5 peak at 2.50 ppm) as internal standard for DMSO-*d*6 solutions. All of the analogs tested in the biological assays have a purity greater than 95% based on LCMS analysis. High resolution mass spectrometry was recorded on Agilent 6210 Time-of-Flight LC/MS system. Confirmation of molecular formulae was accomplished using electrospray ionization in the positive mode with the Agilent Masshunter software (version B.02).

### 978 Synthesis

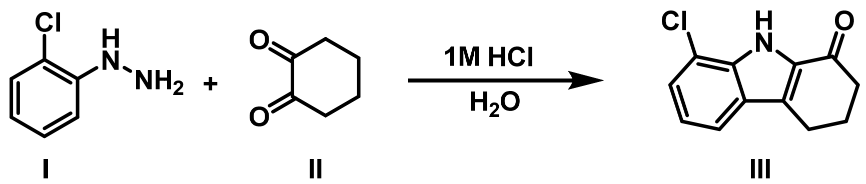

**Method 1** *Step 1:* **8-chloro-2,3,4,9-tetrahydro-1H-carbazol-1-one (III)[2]:** A mixture of a substituted phenylhydrazine **1** (1.0 eq), 1,2-cyclohexanedione **II** (2.0 mmol), and water (50 mL) was stirred at room temperature (23 °C) in a 250 mL round bottom flask. After 16 h, 10 mL of aqueous 1 M HCl was added and the reaction was heated to reflux. After 12 h, the reaction was refrigerated to induce crystallization. The resulting solid was filtered and air dried to provide the crude product. The desired product was purified by flash column chromatography to afford 8-chloro-2,3,4,9-tetrahydro-1H-carbazol-1-one (III).

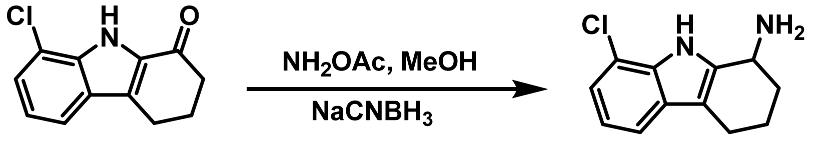


*Step 2:* **8-Chloro-2,3,4,9-tetrahydro-1H-carbazol-1-amine (IV)**: Solid $NH_4OAc$ (5.0 mmol) was added to a stirring solution of 8-chloro-2,3,4,9-tetrahydro-1H-carbazol-1-one **III** (0.5 mmol) dissolved in 20 mL solvent grade $CH_3OH$, and this mixture was allowed to stir at room temperature (23 °C) for 3-6 h. Upon complete consumption of **III**, as determined by thin layer chromatography, solid $NaCNBH_3$ (2.5 mmol) was added and the temperature was raised to 60 °C. After stirring 12-16 h, the reaction was cooled to room temperature (23 °C) and treated with an aqueous 1 M HCl solution. The mixture was extracted with ethyl acetate, and the combined organic layers were washed with brine and dried over anhydrous $Na_2SO_4$, filtered, and concentrated under reduced pressure to afford the crude product. The desired product was purified by flash column chromatography to afford 8-Chloro-2,3,4,9-tetrahydro-1H-carbazol-1-amine (IV). $^1$H NMR (400 MHz, DMSO-$d_6$) δ 10.92 (s, 1H), 7.31 (d, $J$ = 7.8 Hz, 1H), 7.06 (d, $J$ = 7.6 Hz, 1H), 6.92 (t, $J$ = 7.7 Hz, 1H), 3.95 (dd, $J$ = 7.2, 5.5 Hz, 1H), 2.56 (t, $J$ = 6.1 Hz, 2H), 2.26 (s, 2H), 2.11 – 1.86 (m, 1H), 1.77 – 1.59 (m, 1H), 1.53 (dddd, $J$ = 12.4, 9.7, 7.1, 2.1 Hz, 1H).

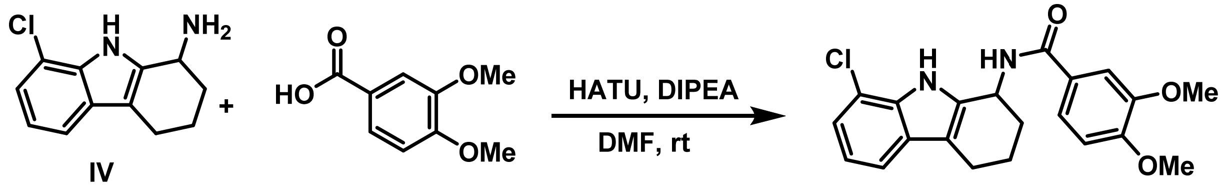

**Method 2 (General Procedure): N-(8-Chloro-2,3,4,9-tetrahydro-1H-carbazol-1-yl)-3,4-dimethoxybenzamide (Table 1: Analog 1):** To a solution of 8-Chloro-2,3,4,9-tetrahydro-1H-carbazol-1-amine (IV) (1.0 eq) in DMF (Volume: 2 ml) was added 2-(3H-[1,2,3]triazolo[4,5-b]pyridin-3-yl)-1,1,3,3-tetramethylisouronium hexafluorophosphate (2.0 eq) and N-ethyl-N-isopropylpropan-2-amine (3.0 eq) and 3,4-dimethoxybenzoic acid (1.0 eq) the reaction mixture was stirred for 12 hrs at room temperature and dilute with water and extract with 3 x 10 mL DCM, washed with brine. The organic layer was dried and concentrated. The crude mixture was diluted with DMSO and purified by reverse phase chromatography (**Method Acidic Standard Gradient**) to afford as a TFA salt, N-(8-chloro-2,3,4,9-tetrahydro-1H-carbazol-1-yl)-3,4-dimethoxybenzamide; LCMS: *m/z* $(M+H)^+$ = 386; $^1$H NMR (400 MHz, dmso) δ 11.15 (s, 1H), 8.64 (d, *J* = 7.5 Hz, 1H), 7.60 – 7.50 (m, 2H), 7.39 (d, *J* = 7.8 Hz, 1H), 7.10 (dd, *J* = 7.6, 1.0 Hz, 1H), 7.01 – 6.92 (m, 2H), 5.31 (q, *J* = 5.3 Hz, 1H), 3.77 (d, *J* = 4.9 Hz, 6H), 2.71 (dt, *J* = 15.4, 5.0 Hz, 2H), 2.66 – 2.56 (m, 2H), 2.03 – 1.85 (m, 2H),

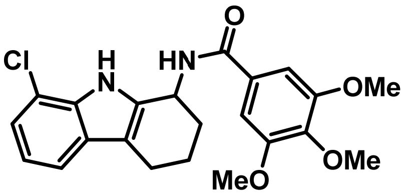


**N-(8-Chloro-2,3,4,9-tetrahydro-1H-carbazol-1-yl)-3,4,5-trimethoxybenzamide (Table 1; Analog 2):** This compound was prepared from **Method 2** using 3,4,5-trimethoxybenzoic acid to afford N-(8-chloro-2,3,4,9-tetrahydro-1H-carbazol-1-yl)-3,4,5-trimethoxybenzamide as a TFA salt. LCMS: *m/z* $(M+H)^+$ = 415; $^1$H NMR (400 MHz, dmso) δ 11.18 (s, 1H), 8.74 (d, *J* = 7.6 Hz, 1H), 7.39 (d, *J* = 7.8 Hz, 1H), 7.28 (s, 2H), 7.11 (dd, *J* = 7.6, 1.0 Hz, 1H), 6.96 (t, *J* = 7.7 Hz, 1H), 5.31 (dd, *J* = 7.5, 4.6 Hz, 1H), 3.79 (s, 6H), 3.68 (s, 3H), 2.72 (dt, *J* = 15.5, 4.9 Hz, 2H), 2.67 – 2.57 (m, 2H), 2.06 – 1.95 (m, 1H), 1.98 – 1.77 (m, 1H).

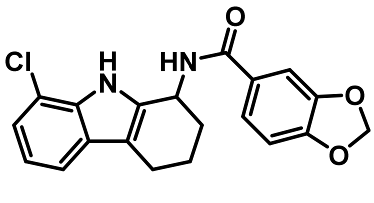


**N-(8-Chloro-2,3,4,9-tetrahydro-1H-carbazol-1-yl)benzo[d][1,3]dioxole-5-carboxamide (Table 1; Analog 3):** This compound was prepared from **Method 2** using benzo[d][1,3]dioxole-

5-carboxylic acid to afford N-(8-chloro-2,3,4,9-tetrahydro-1H-carbazol-1-yl)benzo[d][1,3]dioxole-5-carboxamide as a TFA salt. LCMS: *m/z* $(M+H)^+$ = 369; 1H NMR (400 MHz, dmso) δ 11.13 (s, 1H), 8.60 (d, *J* = 7.5 Hz, 1H), 7.52 (dd, *J* = 8.2, 1.7 Hz, 1H), 7.47 (d, *J* = 1.8 Hz, 1H), 7.39 (d, *J* = 7.8 Hz, 1H), 7.10 (dd, *J* = 7.6, 1.0 Hz, 1H), 7.00 – 6.91 (m, 2H), 6.06 (s, 2H), 5.27 (dd, *J* = 7.4, 4.8 Hz, 1H), 2.65 (dtd, *J* = 21.3, 15.2, 5.3 Hz, 2H), 2.05 – 1.73 (m, 4H).

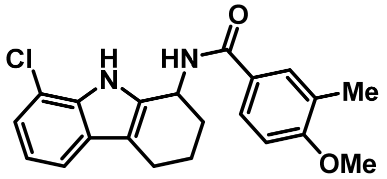


**N-(8-Chloro-2,3,4,9-tetrahydro-1H-carbazol-1-yl)-4-methoxy-3-methylbenzamide (Table 1; Analog 4):** This compound was prepared from **Method 2** using 4-methoxy-3-methylbenzoic acid to afford N-(8-chloro-2,3,4,9-tetrahydro-1H-carbazol-1-yl)-4-methoxy-3-methylbenzamide as a TFA salt. LCMS: *m/z* $(M+H)^+$ = 369; $^1$H NMR (400 MHz, cdcl3) δ 8.85 (s, 1H), 7.68 (d, *J* = 2.2 Hz, 1H), 7.59 (d, *J* = 8.2 Hz, 1H), 7.47 – 7.36 (m, 2H), 7.17 (d, *J* = 7.6 Hz, 1H), 7.02 (t, *J* = 7.8 Hz, 1H), 6.33 (d, *J* = 7.2 Hz, 1H), 5.38 (q, *J* = 5.7 Hz, 1H), 3.79 (s, 3H), 2.85 – 2.67 (m, 2H), 2.44 (s, 3H), 2.31 (td, *J* = 9.7, 5.7 Hz, 2H), 1.99-1.96 (m 2H).

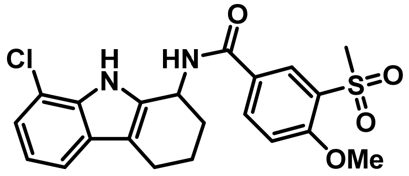


**N-(8-Chloro-2,3,4,9-tetrahydro-1H-carbazol-1-yl)-4-methoxy-3-(methylsulfonyl)benzamide (Table 1; Analog 5):** This compound was prepared from **Method 2** using 4-methoxy-3-(methylsulfonyl)benzoic acid to afford N-(8-Chloro-2,3,4,9-tetrahydro-1H-carbazol-1-yl)-4-methoxy-3-(methylsulfonyl)benzamide as a TFA salt. LCMS: *m/z* $(M+H)^+$ = 432; $^1$H NMR (400 MHz, cdcl3) δ 9.24 (s, 1H), 8.05 (d, *J* = 7.1 Hz, 1H), 7.94 (dd, *J* = 8.3, 6.8 Hz, 1H), 7.90 – 7.81 (m, 2H), 7.33 (d, *J* = 7.8 Hz, 1H), 7.08 (d, *J* = 7.6 Hz, 1H), 6.94 (t, *J* = 7.7 Hz, 1H), 5.36 (q, *J* = 6.1 Hz, 1H), 3.79 (s, 3H), 3.18 (s, 3H), 2.69 (q, *J* = 6.7 Hz, 2H), 2.04 – 1.85 (m, 4H).

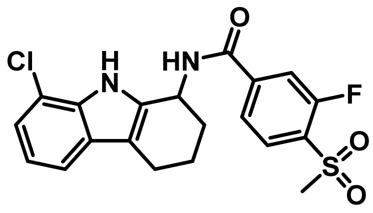


**N-(8-Chloro-2,3,4,9-tetrahydro-1H-carbazol-1-yl)-3-fluoro-4-(methylsulfonyl)benzamide (Table 1; Analog 6):** This compound was prepared from **Method 2** using 3-fluoro-4-(methylsulfonyl)benzoic acid to afford N-(8-chloro-2,3,4,9-tetrahydro-1H-carbazol-1-yl)-3-fluoro-4-(methylsulfonyl)benzamide as a TFA salt. LCMS: *m/z* $(M+H)^+$ = 421; $^1$H NMR (400 MHz, cdcl3) δ 9.24 (s, 1H), 8.05 (d, *J* = 7.1 Hz, 1H), 7.94 (dd, *J* = 8.3, 6.8 Hz, 1H), 7.90 – 7.81 (m, 2H), 7.33 (d, *J* = 7.8 Hz, 1H), 7.08 (d, *J* = 7.6 Hz, 1H), 6.94 (t, *J* = 7.7 Hz, 1H), 5.36 (q, *J* = 6.1 Hz, 1H), 3.18 (s, 3H), 2.69 (q, *J* = 6.7 Hz, 2H), 2.04 – 1.85 (m, 4H).

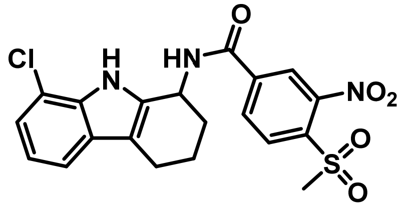


**N-(8-Chloro-2,3,4,9-tetrahydro-1H-carbazol-1-yl)-4-(methylsulfonyl)-3-nitrobenzamide (Table 1; Analog 7):** This compound was prepared from **Method 2** using 4-(methylsulfonyl)-3-nitrobenzoic acid to afford N-(8-chloro-2,3,4,9-tetrahydro-1H-carbazol-1-yl)-4-(methylsulfonyl)-3-nitrobenzamide as a TFA salt. LCMS: *m/z* $(M+H)^+$ = 448; 1H NMR (400 MHz, cdcl3) δ 8.32 (d, *J* = 1.6 Hz, 1H), 8.26 – 8.15 (m, 2H), 7.34 (dd, *J* = 7.8, 1.0 Hz, 1H), 7.10 (dd, *J* = 7.7, 1.0 Hz, 1H), 6.95 (t, *J* = 7.7 Hz, 1H), 5.35 (t, *J* = 5.3 Hz, 1H), 4.68 (s, 1H), 3.39 (s, 3H), 2.79 – 2.60 (m, 2H), 2.20 (ddt, *J* = 12.9, 8.7, 4.2 Hz, 2H), 2.04 – 1.87 (m, 2H).

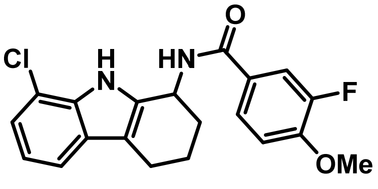


**N-(8-chloro-2,3,4,9-tetrahydro-1H-carbazol-1-yl)-3-fluoro-4-methoxybenzamide (Table 1; Analog 8):** This compound was prepared from **Method 2** using 3-fluoro-4-methoxybenzoic acid to afford N-(8-chloro-2,3,4,9-tetrahydro-1H-carbazol-1-yl)-3-fluoro-4-methoxybenzamide as a

TFA salt. LCMS: *m/z* $(M+H)^+$ = 373; $^1H$ NMR (400 MHz, cdcl3) δ 7.59 (dd, *J* = 8.2, 2.1 Hz, 1H), 7.35 (dd, *J* = 7.8, 1.0 Hz, 1H), 7.28 – 7.20 (m, 1H), 7.12 – 6.98 (m, 2H), 6.94 (t, *J* = 7.7 Hz, 1H), 5.32 (t, *J* = 5.2 Hz, 1H), 3.88 (s, 3H), 2.77 – 2.60 (m, 2H), 2.25 – 2.11 (m, 2H), 1.94 (dq, *J* = 14.0, 5.4 Hz, 2H).

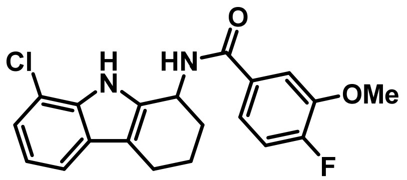


**N-(8-Chloro-2,3,4,9-tetrahydro-1H-carbazol-1-yl)-4-fluoro-3-methoxybenzamide (Table 1; Analog 9):** This compound was prepared from **Method 2** using 4-fluoro-3-methoxybenzoic acid to afford N-(8-chloro-2,3,4,9-tetrahydro-1H-carbazol-1-yl)-4-fluoro-3-methoxybenzamide as a TFA salt. LCMS: *m/z* $(M+H)^+$ = 373; $^1H$ NMR (400 MHz, cdcl3) δ 7.49 (dd, *J* = 8.2, 2.1 Hz, 1H), 7.34 (dd, *J* = 7.8, 1.0 Hz, 1H), 7.29 – 7.21 (m, 1H), 7.13 – 6.99 (m, 2H), 6.95 (t, *J* = 7.7 Hz, 1H), 5.33 (t, *J* = 5.2 Hz, 1H), 3.89 (s, 3H), 2.79 – 2.60 (m, 2H), 2.26 – 2.12 (m, 2H), 1.95 (dq, *J* = 14.0, 5.4 Hz, 2H).

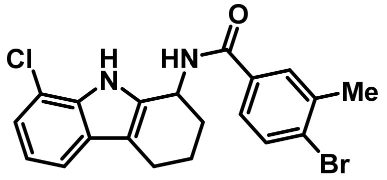


**4-Bromo-N-(8-chloro-2,3,4,9-tetrahydro-1H-carbazol-1-yl)-3-methylbenzamide (Table 1; Analog 10):** This compound was prepared from **Method 2** using 4-bromo-3-methylbenzoic acid to afford 4-bromo-N-(8-chloro-2,3,4,9-tetrahydro-1H-carbazol-1-yl)-3-methylbenzamide as a TFA salt. LCMS: *m/z* $(M+H)^+$ = 419; $^1H$ NMR (400 MHz, cdcl3) δ 8.85 (s, 1H), 7.68 (d, *J* = 2.2 Hz, 1H), 7.59 (d, *J* = 8.2 Hz, 1H), 7.47 – 7.36 (m, 2H), 7.17 (d, *J* = 7.6 Hz, 1H), 7.02 (t, *J* = 7.8 Hz, 1H), 6.33 (d, *J* = 7.2 Hz, 1H), 5.38 (q, *J* = 5.7 Hz, 1H), 2.85 – 2.67 (m, 2H), 2.44 (s, 3H), 2.31 (td, *J* = 9.7, 5.7 Hz, 2H), 1.99-1.96 (m 2H).

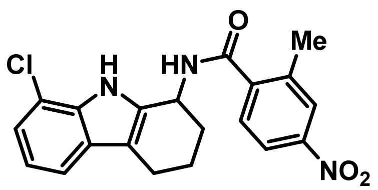

**N-(8-Chloro-2,3,4,9-tetrahydro-1H-carbazol-1-yl)-2-methyl-4-nitrobenzamide (Table 1; Analog 11):** This compound was prepared from **Method 2** using 2-methyl-4-nitrobenzoic acidto afford N-(8-chloro-2,3,4,9-tetrahydro-1H-carbazol-1-yl)-2-methyl-4-nitrobenzamide as a TFA salt. LCMS: *m/z* $(M+H)^+$ = 384; $^1$H NMR (400 MHz, cdcl3) δ 9.66 (d, *J* = 7.1 Hz, 1H), 8.02 – 7.72 (m, 1H), 7.45 (t, *J* = 8.5 Hz, 1H), 7.27 – 7.15 (m, 2H), 7.03 – 6.71 (m, 2H), 5.18 (d, *J* = 7.8 Hz, 1H), 4.54 (s, 1H), 2.53 (s, 3H), 2.42 – 2.28 (m, 4H), 1.79 (d, *J* = 28.9 Hz, 2H).

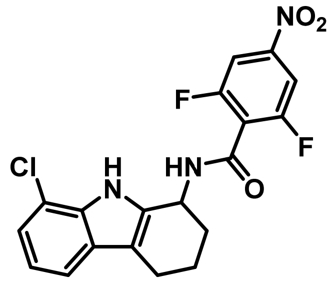


**N-(8-chloro-2,3,4,9-tetrahydro-1H-carbazol-1-yl)-2,6-difluoro-4-nitrobenzamide (Table 1; Analog 12):** This compound was prepared from **Method 2** using 2,6-difluoro-4-nitrobenzoic acid to afford N-(8-chloro-2,3,4,9-tetrahydro-1H-carbazol-1-yl)-2,6-difluoro-4-nitrobenzamide as a TFA salt. LCMS: *m/z* $(M+H)^+$ = 406; $^1$H NMR (400 MHz, cdcl3) δ 8.66 (s, 1H), 7.58 – 7.50 (m, 2H), 7.41 (d, *J* = 7.9 Hz, 1H), 7.19 (d, *J* = 7.6 Hz, 1H), 7.04 (t, *J* = 7.8 Hz, 1H), 6.39 (d, *J* = 7.2 Hz, 1H), 5.39 (q, *J* = 5.7 Hz, 1H), 2.86 – 2.67 (m, 2H), 2.31 (td, *J* = 10.7, 5.7 Hz, 2H), 2.00 (ddt, *J* = 30.7, 10.3, 5.9 Hz, 2H).

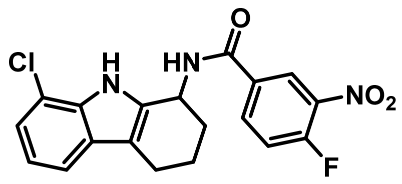


**N-(8-chloro-2,3,4,9-tetrahydro-1H-carbazol-1-yl)-4-fluoro-3-nitrobenzamide (Table 1; Analog 13):** This compound was prepared from **Method 2** using 4-fluoro-3-nitrobenzoic acid to afford N-(8-chloro-2,3,4,9-tetrahydro-1H-carbazol-1-yl)-4-fluoro-3-nitrobenzamide as a TFA salt. LCMS: *m/z* $(M+H)^+$ = 388; 1H NMR (400 MHz, cdcl3) δ 8.65 – 8.60 (m, 1H), 7.92 – 7.84 (m, 2H), 7.79 (dd, *J* = 8.2, 2.2 Hz, 1H), 7.74 – 7.67 (m, 1H), 7.58 – 7.50 (m, 2H), 5.39 (q, *J* = 5.7 Hz, 1H), 2.84-2.65 (m, 4H), 2.56-2.30 (m, 2H).

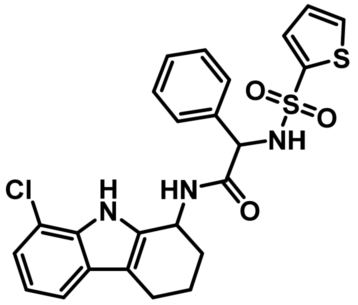


**N-(8-Chloro-2,3,4,9-tetrahydro-1H-carbazol-1-yl)-2-phenyl-2-(thiophene-2-sulfonamido)acetamide (Table 1; Analog 14):** This compound was prepared from **Method 2** using 2-phenyl-2-(thiophene-2-sulfonamido)acetic acid to afford N-(8-chloro-2,3,4,9-tetrahydro-1H-carbazol-1-yl)-2-phenyl-2-(thiophene-2-sulfonamido)acetamide as a TFA salt. LCMS: *m/z* $(M+H)^+$ = 500.1; $^1$H NMR (400 MHz, dmso) δ 10.89 (s, 1H), 10.72 (s, 1H), 8.80 (dd, *J* = 20.1, 9.4 Hz, 1H), 8.67 (d, *J* = 7.1 Hz, 1H), 7.85 (dd, *J* = 5.0, 1.4 Hz, 1H), 7.61 (dd, *J* = 5.0, 1.4 Hz, 1H), 7.48 (dd, *J* = 3.7, 1.4 Hz, 1H), 7.43 – 7.30 (m, 2H), 7.30 – 7.05 (m, 2H), 6.96 (dt, *J* = 12.7, 7.7 Hz, 2H), 6.86 (dd, *J* = 5.0, 3.7 Hz, 1H), 4.88 – 4.81 (m, 1H), 2.67 (t, *J* = 17.0 Hz, 2H), 2.52 (d, *J* = 7.1 Hz, 2H), 1.71 (dd, *J* = 17.1, 6.9 Hz, 2H).

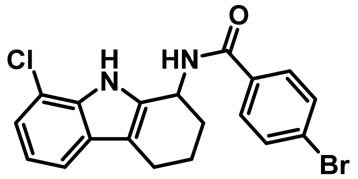


**4-Bromo-N-(8-chloro-2,3,4,9-tetrahydro-1H-carbazol-1-yl)benzamide (Table 1; Analog 15):** This compound was prepared from **Method 2** using 4-bromobenzoic acid to afford 4-bromo-N-(8-chloro-2,3,4,9-tetrahydro-1H-carbazol-1-yl)benzamide as a TFA salt. LCMS: *m/z* $(M+H)^+$ = 404; $^1$H NMR (400 MHz, dmso) δ 11.17 (s, 1H), 8.86 (d, *J* = 7.6 Hz, 1H), 7.90 – 7.82 (m, 2H), 7.68 – 7.60 (m, 2H), 7.39 (d, *J* = 7.8 Hz, 1H), 7.10 (dd, *J* = 7.6, 1.0 Hz, 1H), 6.96 (t, *J* = 7.7 Hz, 1H), 5.33 – 5.25 (m, 1H), 2.74 – 2.57 (m, 2H), 2.04 – 1.75 (m, 4H).

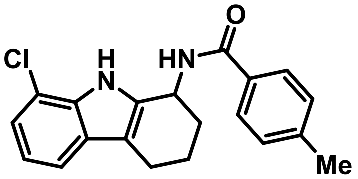

**N-(8-chloro-2,3,4,9-tetrahydro-1H-carbazol-1-yl)-4-methylbenzamide (Table 1; Analog 16):** This compound was prepared from **Method 2** using 4-methylbenzoic acid to afford N-(8-chloro-2,3,4,9-tetrahydro-1H-carbazol-1-yl)-4-methylbenzamide as a TFA salt. LCMS: *m/z* $(M+H)^+$ = 339; $^1$H NMR (400 MHz, dmso) δ 11.13 (s, 1H), 8.67 (d, *J* = 7.6 Hz, 1H), 7.85 – 7.79 (m, 2H), 7.39 (d, *J* = 7.8 Hz, 1H), 7.22 (d, *J* = 8.0 Hz, 2H), 7.10 (dd, *J* = 7.6, 0.9 Hz, 1H), 6.96 (t, *J* = 7.7 Hz, 1H), 5.30 (d, *J* = 6.7 Hz, 1H), 2.65 – 2.56 (m, 2H), 2.33 (s, 3H), 2.00 – 1.87 (m, 2H), 1.79 (d, *J* = 12.2 Hz, 2H).

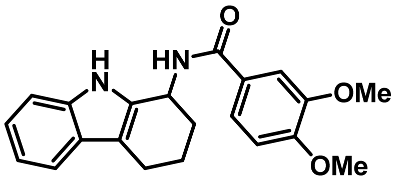


**3,4-Dimethoxy-N-(2,3,4,9-tetrahydro-1H-carbazol-1-yl)benzamide (Table 2; Analog 17):** This compound was prepared from **Method 2** using 2,3,4,9-tetrahydro-1H-carbazol-1-amine and 3,4-dimethoxybenzoic acid to afford 3,4-dimethoxy-N-(2,3,4,9-tetrahydro-1H-carbazol-1-yl)benzamide as a TFA salt. LCMS: *m/z* $(M+H)^+$ = 351; $^1$H NMR (400 MHz, dmso) δ 10.71 (s, 1H), 8.64 (d, *J* = 8.1 Hz, 1H), 7.61 – 7.51 (m, 2H), 7.51 – 7.35 (m, 1H), 7.30 – 7.24 (m, 1H), 7.05 – 6.94 (m, 2H), 6.93 (ddd, *J* = 7.9, 7.1, 1.1 Hz, 1H), 5.33 (q, *J* = 5.1 Hz, 1H), 3.78 (d, *J* = 5.0 Hz, 6H), 2.66 (dt, *J* = 6.5, 3.2 Hz, 2H), 2.02 (tt, *J* = 8.8, 4.3 Hz, 2H), 1.92 – 1.75 (m, 2H).

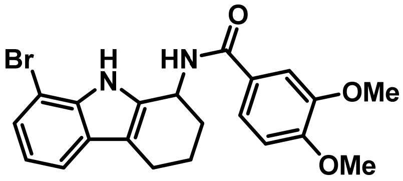


**N-(8-Bromo-2,3,4,9-tetrahydro-1H-carbazol-1-yl)-3,4-dimethoxybenzamide (Table 2; Analog 18):** This compound was prepared from **Method 2** using 8-bromo-2,3,4,9-tetrahydro-1H-carbazol-1-amine and 3,4-dimethoxybenzoic acid to afford N-(8-bromo-2,3,4,9-tetrahydro-1H-carbazol-1-yl)-3,4-dimethoxybenzamide as a TFA salt. LCMS: *m/z* $(M+H)^+$ = 429; $^1$H NMR (400 MHz, cdcl3) δ 8.85 (s, 1H), 7.48 – 7.41 (m, 2H), 7.35 – 7.24 (m, 2H), 6.97 (t, *J* = 7.7 Hz, 1H), 6.85 (d, *J* = 8.4 Hz, 1H), 6.31 (d, *J* = 7.2 Hz, 1H), 5.39 (q, *J* = 5.8 Hz, 1H), 3.94 (d, *J* = 14.0 Hz, 6H), 2.85 – 2.67 (m, 2H), 2.31 (td, *J* = 9.2, 5.6 Hz, 2H), 1.99 (tt, *J* = 6.7, 4.1 Hz, 2H).

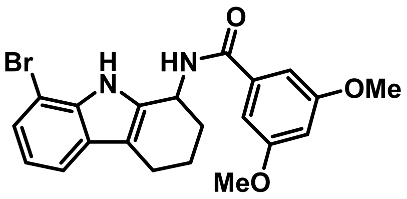


**N-(8-bromo-2,3,4,9-tetrahydro-1H-carbazol-1-yl)-3,5-dimethoxybenzamide (Table 2; Analog 19):** This compound was prepared from **Method 2** using 8-bromo-2,3,4,9-tetrahydro-1H-carbazol-1-amine and 3,5-dimethoxybenzoic acid to afford N-(8-bromo-2,3,4,9-tetrahydro-1H-carbazol-1-yl)-3,5-dimethoxybenzamide as a TFA salt. LCMS: *m/z* $(M+H)^+$ = 429; $^1$H NMR (400 MHz, dmso) δ 11.01 (s, 1H), 8.74 (d, *J* = 7.4 Hz, 1H), 7.43 (d, *J* = 7.8 Hz, 1H), 7.25 (d, *J* = 7.6 Hz, 1H), 7.09 (d, *J* = 2.3 Hz, 2H), 6.91 (t, *J* = 7.7 Hz, 1H), 6.60 (t, *J* = 2.3 Hz, 1H), 5.33 – 5.25 (m, 1H), 3.75 (s, 6H), 2.75 – 2.65 (m, 2H), 2.60 (d, *J* = 15.5 Hz, 2H), 2.02 – 1.85 (m, 2H).

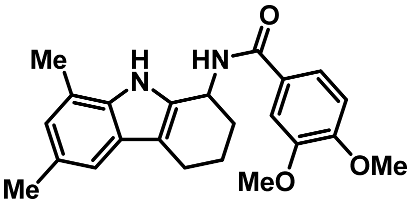


**N-(6,8-Dimethyl-2,3,4,9-tetrahydro-1H-carbazol-1-yl)-3,4-dimethoxybenzamide (Table 1; Analog 20):** This compound was prepared from **Method 2** using 5,8-dimethyl-2,3,4,9-tetrahydro-1H-carbazol-1-amine and 3,4-dimethoxybenzoic acid to afford N-(6,8-dimethyl-2,3,4,9-tetrahydro-1H-carbazol-1-yl)-3,4-dimethoxybenzamide as a TFA salt. LCMS: *m/z* $(M+H)^+$ = 379; $^1$H NMR (400 MHz, dmso) δ 10.52 (s, 1H), 8.61 (d, *J* = 7.8 Hz, 1H), 7.82 (s, 1H), 7.64 – 7.51 (m, 1H), 7.51 – 7.41 (m, 1H), 6.65 (s, 1H), 5.33 – 5.25 (m, 1H), 3.77 (dd, *J* = 4.5, 1.8 Hz, 6H), 2.65-2.63 (m, 2H), 2.62 – 2.49 (m, 2H), 2.34 (s, 3H), 2.30 (s, 3H), 1.90 – 1.76 (m, 2H).

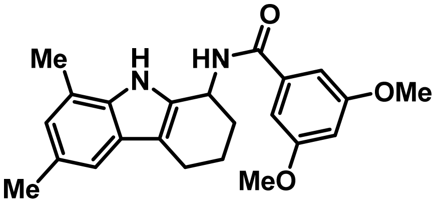


**N-(6,8-Dimethyl-2,3,4,9-tetrahydro-1H-carbazol-1-yl)-3,5-dimethoxybenzamide (Table 2; Analog 21):** This compound was prepared from **Method 2** using 5,8-dimethyl-2,3,4,9-tetrahydro-1H-carbazol-1-amine and 3,5-dimethoxybenzoic acid to afford N-(6,8-dimethyl-2,3,4,9-

tetrahydro-1H-carbazol-1-yl)-3,5-dimethoxybenzamide as a TFA salt. LCMS: $m/z$ $(M+H)^+ = 379$; $^1H$ NMR (400 MHz, cdcl3) δ 9.55 (s, 1H), 8.34 (dd, $J$ = 28.8, 4.7 Hz, 1H), 8.05 (dd, $J$ = 8.8, 2.3 Hz, 1H), 7.17 (d, $J$ = 7.8 Hz, 1H), 6.93 (dd, $J$ = 26.5, 8.2 Hz, 1H), 6.77 (t, $J$ = 7.7 Hz, 1H), 5.20 (d, $J$ = 6.5 Hz, 1H), 3.80 (s, 6H), 2.53 (d, $J$ = 8.0 Hz, 2H), 2.38 (s, 6H), 2.04 – 1.61 (m, 4H).

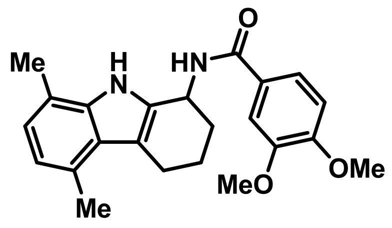


**N-(5,8-dimethyl-2,3,4,9-tetrahydro-1H-carbazol-1-yl)-3,4-dimethoxybenzamide (Table 1; Analog 22):** This compound was prepared from **Method 2** using 5,8-dimethyl-2,3,4,9-tetrahydro-1H-carbazol-1-amine and 3,4-dimethoxybenzoic acid to afford N-(5,8-dimethyl-2,3,4,9-tetrahydro-1H-carbazol-1-yl)-3,4-dimethoxybenzamide as a TFA salt. LCMS: $m/z$ $(M+H)^+ = 379$; $^1H$ NMR (400 MHz, dmso) δ 10.57 (s, 1H), 8.59 (d, $J$ = 7.7 Hz, 1H), 7.60 – 7.50 (m, 1H), 6.97 (d, $J$ = 8.4 Hz, 1H), 6.66 (d, $J$ = 7.1 Hz, 1H), 6.55 (d, $J$ = 7.1 Hz, 1H), 5.29 (d, $J$ = 6.7 Hz, 1H), 3.77 (d, $J$ = 5.0 Hz, 6H), 2.88 (dd, $J$ = 15.7, 7.0 Hz, 2H), 2.52 (s, 3H), 2.32 (s, 3H), 1.94 (d, $J$ = 7.1 Hz, 2H), 1.82 (ddd, $J$ = 18.2, 9.0, 3.5 Hz, 2H).

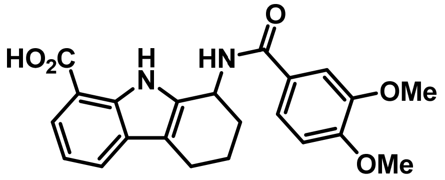


**1-(3,4-Dimethoxybenzamido)-2,3,4,9-tetrahydro-1H-carbazole-8-carboxylic acid (Table 2; Analog 23):** This compound was prepared from **Method 2** using 1-amino-2,3,4,9-tetrahydro-1H-carbazole-8-carboxylic acid and 3,4-dimethoxybenzoic acid to afford 1-(3,4-dimethoxybenzamido)-2,3,4,9-tetrahydro-1H-carbazole-8-carboxylic acid as a TFA salt. LCMS: $m/z$ $(M+H)^+ = 395$; $^1H$ NMR (400 MHz, dmso) δ 10.57 (s, 1H), 8.59 (d, $J$ = 7.7 Hz, 1H), 7.60 – 7.50 (m, 2H), 6.97 (d, $J$ = 8.4 Hz, 1H), 6.66 (d, $J$ = 7.1 Hz, 1H), 6.55 (d, $J$ = 7.1 Hz, 1H), 5.29 (d, $J$ = 6.7 Hz, 1H), 3.85 (d, $J$ = 4.0 Hz, 3H), 3.77 (d, $J$ = 5.0 Hz, 3H), 2.88 (dd, $J$ = 15.7, 7.0 Hz, 2H), 1.94 (d, $J$ = 7.1 Hz, 2H), 1.82 (ddd, $J$ = 18.2, 9.0, 3.5 Hz, 2H).

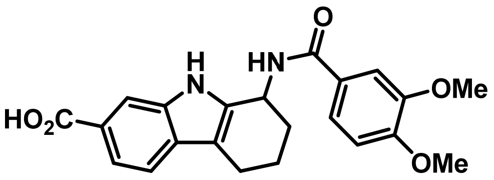


**1-(3,4-Dimethoxybenzamido)-2,3,4,9-tetrahydro-1H-carbazole-7-carboxylic acid (Table 2; Analog 24):** This compound was prepared from **Method 2** using 1-amino-2,3,4,9-tetrahydro-1H-carbazole-7-carboxylic acid and 3,4-dimethoxybenzoic acid to afford 1-(3,4-dimethoxybenzamido)-2,3,4,9-tetrahydro-1H-carbazole-7-carboxylic acid as a TFA salt. LCMS: *m/z* $(M+H)^+$ = 395; $^1$H NMR (400 MHz, cdcl3) δ 9.29 (s, 1H), 7.78 (d, *J* = 7.5 Hz, 1H), 7.54 (d, *J* = 8.1 Hz, 1H), 7.44 (d, *J* = 2.0 Hz, 1H), 7.29 (d, *J* = 2.1 Hz, 1H), 7.17 (t, *J* = 7.8 Hz, 1H), 6.87 (d, *J* = 8.4 Hz, 1H), 6.35 (d, *J* = 6.8 Hz, 1H), 5.37 (d, *J* = 6.3 Hz, 1H), 3.85 (s, 6H), 3.00 (d, *J* = 17.1 Hz, 2H), 2.38 – 2.30 (m, 2H), 1.96-1.94 (m, 2H).

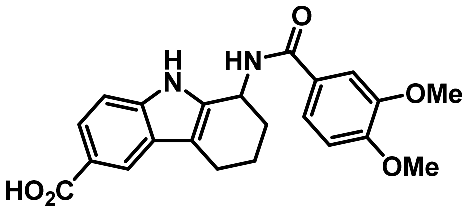


**1-(3,4-Dimethoxybenzamido)-2,3,4,9-tetrahydro-1H-carbazole-6-carboxylic acid (Table 2; Analog 25):** This compound was prepared from **Method 2** using 1-amino-2,3,4,9-tetrahydro-1H-carbazole-6-carboxylic acid and 3,4-dimethoxybenzoic acid to afford 1-(3,4-dimethoxybenzamido)-2,3,4,9-tetrahydro-1H-carbazole-6-carboxylic acid as a TFA salt. LCMS: *m/z* $(M+H)^+$ = 395; $^1$H NMR (400 MHz, cdcl3) δ 8.20 – 8.03 (m, 1H), 7.60 – 7.46 (m, 3H), 7.26 (s, 1H), 6.84 (dd, *J* = 8.2, 2.7 Hz, 1H), 5.38 (d, *J* = 6.8 Hz, 1H), 3.91 (s, 6H), 2.08 – 1.81 (m, 4H), 1.86-1.84 (m, 2H).

### 527 Synthesis

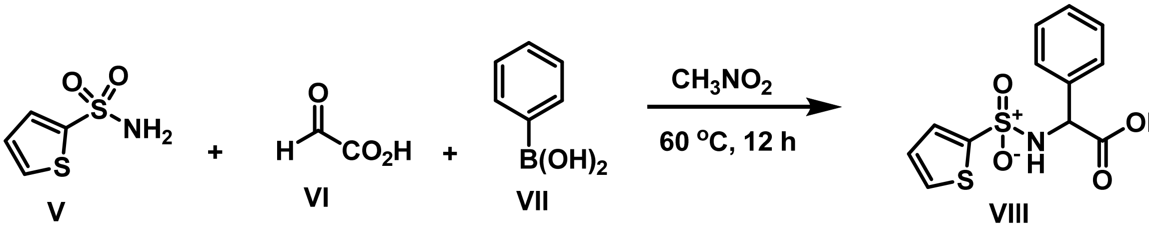


**Method 3 (Petasis-Borono Mannich Reaction): 2-Phenyl-2-(thiophene-2-sulfonamido)acetic acid (VIII)[1]:** A 5–mL microwave tube was charged with a magnetic stirring bar, sulfonamide **V** (1.0 equiv), glyoxylic acid monohydrate **VI** (1.3 equiv), boronic acid **VII** (2.0 equiv), and nitromethane (1.5 mL, 0.17 M wrt sulfonamide) and firmly closed with cap. The resulting mixture was stirred at 60 °C for 12 h. After cooling to r.t., the mixture was diluted with acetone and filtered through a short plug of Celite/silica gel. The plug was rinsed with additional acetone and the filtrate was concentrated under reduced pressure. Purification of the crude residue by flash column chromatography afforded the product 2-Phenyl-2-(thiophene-2-sulfonamido)acetic acid (VIII).

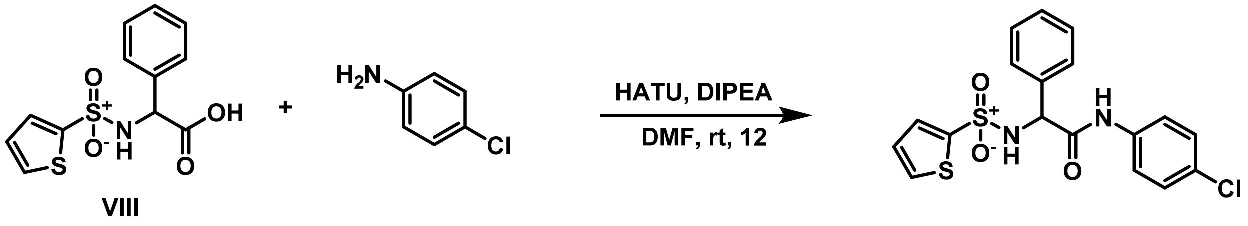


**Method 4 (General Procedure): N-(4-Chlorophenyl)-2-phenyl-2-(thiophene-2-sulfonamido)acetamide (Table 5: Analog 28):** To a solution of **2**-Phenyl-2-(thiophene-2-sulfonamido)acetic acid (1.0 eq) in DMF (Volume: 2 ml) was added 2-(3H-[1,2,3]triazolo[4,5-b]pyridin-3-yl)-1,1,3,3-tetramethylisouronium hexafluorophosphate (2.0 eq) and N-ethyl-N-isopropylpropan-2-amine (3.0 eq) and 4-chloroaniline (1.0 eq) the reaction mixture was stirred for 12 hrs at room temperature and dilute with water and extract with 3 x 10 mL DCM, washed with brine. The organic layer was dried and concentrated. The crude mixture was diluted with DMSO and purified by reverse phase chromatography (**Method Acidic Standard Gradient**) to afford as a TFA salt, N-(4-chlorophenyl)-2-phenyl-2-(thiophene-2-sulfonamido)acetamide; LCMS: *m/z* $(M+H)^+$ = 407; $^1$H NMR (400 MHz, dmso) δ 10.41 (s, 1H), 9.01 (d, *J* = 9.4 Hz, 1H), 7.77 (dd, *J* = 5.0, 1.4 Hz, 1H), 7.49 – 7.20 (m, 9H), 6.98 (dd, *J* = 5.0, 3.7 Hz, 1H), 5.20 (d, *J* = 9.4 Hz, 1H).

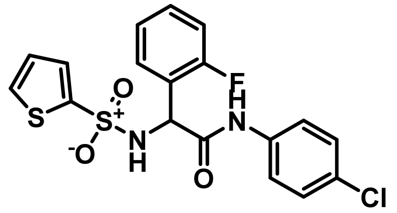


**N-(4-Chlorophenyl)-2-(2-fluorophenyl)-2-(thiophene-2-sulfonamido)acetamide (Table 3; Analog 29):** This compound was prepared from **Method 4** using 4-chloroaniline and 2-(2-fluorophenyl)-2-(thiophene-2-sulfonamido)acetic acid ( This compound was prepared from **Method 3**) to afford N-(4-chlorophenyl)-2-(2-fluorophenyl)-2-(thiophene-2-sulfonamido)acetamide as a TFA salt. LCMS: *m/z* $(M+H)^+$ = 425; $^1H$ NMR (400 MHz, dmso) δ 10.40 (s, 1H), 8.99 (d, *J* = 9.1 Hz, 1H), 7.81 – 7.68 (m, 1H), 7.53 – 7.46 (m, 1H), 7.50 – 7.39 (m, 2H), 7.42 – 7.27 (m, 2H), 7.18 – 7.05 (m, 2H), 6.99 (dd, *J* = 5.0, 3.7 Hz, 1H), 5.44 (d, *J* = 9.1 Hz, 1H).

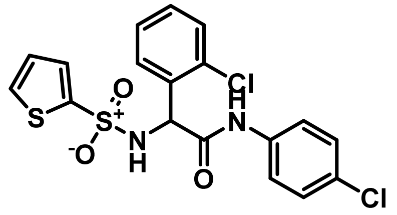


**2-(2-Chlorophenyl)-N-(4-chlorophenyl)-2-(thiophene-2-sulfonamido)acetamide (Table 3; Analog 30):** This compound was prepared from **Method 4** using 4-chloroaniline and 2-(2-chlorophenyl)-2-(thiophene-2-sulfonamido)acetic acid (This compound was prepared from **Method 3**) to afford 2-(2-chlorophenyl)-N-(4-chlorophenyl)-2-(thiophene-2-sulfonamido)acetamide as a TFA salt. LCMS: *m/z* $(M+H)^+$ = 441; $^1H$ NMR (400 MHz, dmso) δ 10.39 (s, 1H), 8.96 (d, *J* = 8.9 Hz, 1H), 7.78 (dd, *J* = 5.0, 1.4 Hz, 1H), 7.57 – 7.42 (m, 2H), 7.40 (dd, *J* = 7.9, 1.5 Hz, 1H), 7.37 – 7.29 (m, 2H), 7.28 (td, *J* = 7.6, 1.9 Hz, 1H), 7.23 (td, *J* = 7.5, 1.5 Hz, 1H), 6.99 (dd, *J* = 5.0, 3.7 Hz, 1H), 5.51 (d, *J* = 8.9 Hz, 1H).

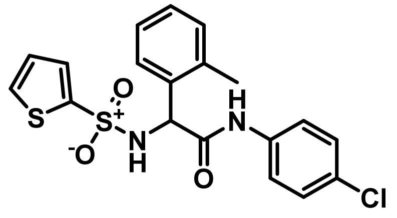

**N-(4-Chlorophenyl)-2-(thiophene-2-sulfonamido)-2-(o-tolyl)acetamide (Table 3; Analog 31):** This compound was prepared from **Method 4** using 4-chloroaniline and 2-(thiophene-2-sulfonamido)-2-(o-tolyl)acetic acid (This compound was prepared from **Method 3**) to afford 2-(2-chlorophenyl)-N-(4-chlorophenyl)-2-(thiophene-2-sulfonamido)acetamide as a TFA salt. LCMS: *m/z* $(M+H)^+$ = 421; $^1H$ NMR (400 MHz, dmso) δ 10.20 (s, 1H), 8.69 (d, *J* = 8.8 Hz, 1H), 7.79 (ddd, *J* = 12.7, 5.0, 1.4 Hz, 1H), 7.53 – 7.40 (m, 2H), 7.36 – 7.23 (m, 2H), 7.22 – 7.14 (m, 2H), 7.14 – 6.96 (m, 2H), 5.26 (d, *J* = 8.8 Hz, 1H), 2.32 (s, 3H).

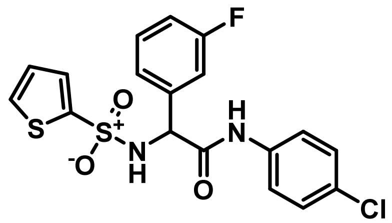

**N-(4-Chlorophenyl)-2-(3-fluorophenyl)-2-(thiophene-2-sulfonamido)acetamide (Table 3; Analog 32):** This compound was prepared from **Method 4** using 4-chloroaniline and 2-(3-fluorophenyl)-2-(thiophene-2-sulfonamido)acetic acid (This compound was prepared from **Method 3**) to afford 2-(2-chlorophenyl)-N-(4-chlorophenyl)-2-(thiophene-2-sulfonamido)acetamide as a TFA salt. LCMS: *m/z* $(M+H)^+$ = 425; $^1H$ NMR (400 MHz, dmso) δ 10.47 (s, 1H), 9.10 (d, *J* = 9.6 Hz, 1H), 7.77 (dd, *J* = 5.0, 1.4 Hz, 1H), 7.49 – 7.39 (m, 2H), 7.37 – 7.27 (m, 2H), 7.27 – 7.17 (m, 2H), 7.14 – 7.03 (m, 1H), 6.98 (dd, *J* = 5.0, 3.7 Hz, 1H), 5.25 (d, *J* = 9.6 Hz, 1H).

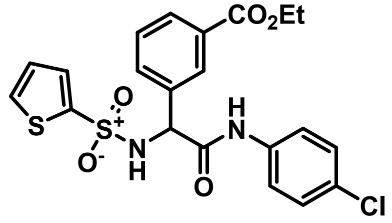

**Ethyl 3-(2-((4-chlorophenyl)amino)-2-oxo-1-(thiophene-2-sulfonamido)ethyl)benzoate (Table 3; Analog 33):** This compound was prepared from **Method 4** using 4-chloroaniline and 2-

(thiophene-2-sulfonamido)-2-(o-tolyl)acetic acid (This compound was prepared from **Method 3**) to afford 2-(2-chlorophenyl)-N-(4-chlorophenyl)-2-(thiophene-2-sulfonamido)acetamide as a TFA salt. LCMS: *m/z* $(M+H)^+$ = 479; 1H NMR (400 MHz, dmso) δ 10.48 (s, 1H), 9.18 (d, *J* = 9.4 Hz, 1H), 8.01 (t, *J* = 1.8 Hz, 1H), 7.82 (dt, *J* = 7.9, 1.4 Hz, 1H), 7.76 (dd, *J* = 5.0, 1.4 Hz, 1H), 7.66 (d, *J* = 7.8 Hz, 1H), 7.48 – 7.39 (m, 4H), 7.36 – 7.28 (m, 2H), 6.96 (dd, *J* = 5.0, 3.7 Hz, 1H), 5.29 (d, *J* = 9.4 Hz, 1H), 4.30 (q, *J* = 7.1 Hz, 2H), 1.31 (t, *J* = 7.1 Hz, 3H).

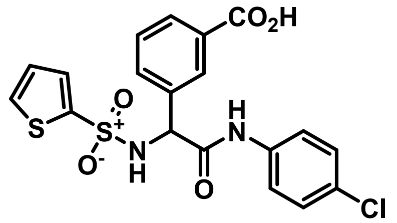


**3-(2-((4-Chlorophenyl)amino)-2-oxo-1-(thiophene-2-sulfonamido)ethyl)benzoic acid (Table 3; Analog 34):** This compound was prepared from **Method 4** using 4-chloroaniline and 2-(thiophene-2-sulfonamido)-2-(o-tolyl)acetic acid (This compound was prepared from **Method 3**) to afford 2-(2-chlorophenyl)-N-(4-chlorophenyl)-2-(thiophene-2-sulfonamido)acetamide as a TFA salt. LCMS: *m/z* $(M+H)^+$ = 451; $^1$H NMR (400 MHz, dmso) δ 13.00 (s, 1H), 10.46 (s, 1H), 9.15 (d, *J* = 9.3 Hz, 1H), 8.02 (t, *J* = 1.8 Hz, 1H), 7.85 – 7.73 (m, 2H), 7.62 (dt, *J* = 7.8, 1.5 Hz, 1H), 7.53 – 7.37 (m, 2H), 7.36 – 7.28 (m, 2H), 6.97 (dd, *J* = 5.0, 3.7 Hz, 1H), 5.28 (d, *J* = 9.4 Hz, 1H).

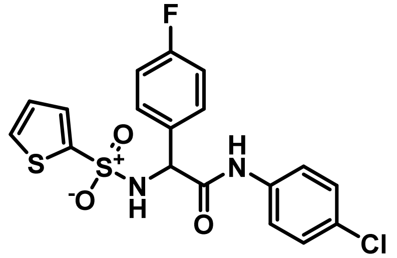


**N-(4-Chlorophenyl)-2-(4-fluorophenyl)-2-(thiophene-2-sulfonamido)acetamide (Table 3; Analog 35):** This compound was prepared from **Method 4** using 4-chloroaniline and 2-(4-fluorophenyl)-2-(thiophene-2-sulfonamido)acetic acid (This compound was prepared from **Method 3**) to afford N-(4-chlorophenyl)-2-(4-fluorophenyl)-2-(thiophene-2-sulfonamido)acetamide as a TFA salt. LCMS: *m/z* $(M+H)^+$ = 425; $^1$H NMR (400 MHz, dmso) δ

10.42 (s, 1H), 9.05 (d, *J* = 9.5 Hz, 1H), 7.77 (dd, *J* = 5.0, 1.4 Hz, 1H), 7.49 – 7.38 (m, 4H), 7.36 – 7.27 (m, 2H), 7.18 – 7.05 (m, 2H), 6.98 (dd, *J* = 5.0, 3.7 Hz, 1H), 5.21 (d, *J* = 9.5 Hz, 1H).

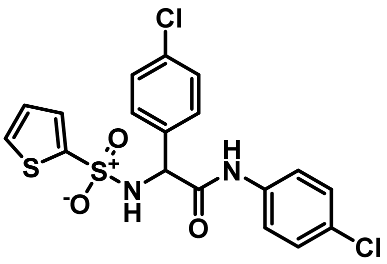


**N,2-bis(4-Chlorophenyl)-2-(thiophene-2-sulfonamido)acetamide (Table 3; Analog 36):** This compound was prepared from **Method 4** using 4-chloroaniline and 2-(4-chlorophenyl)-2-(thiophene-2-sulfonamido)acetic acid (This compound was prepared from **Method 3**) to afford N,2-bis(4-chlorophenyl)-2-(thiophene-2-sulfonamido)acetamide as a TFA salt. LCMS: *m/z* $(M+H)^+$ = 441; $^1$H NMR (400 MHz, dmso) δ 10.44 (s, 1H), 9.09 (d, *J* = 9.5 Hz, 1H), 7.78 (dd, *J* = 5.0, 1.4 Hz, 1H), 7.48 – 7.28 (m, 8H), 6.99 (dd, *J* = 5.0, 3.7 Hz, 1H), 5.22 (d, *J* = 9.4 Hz, 1H).

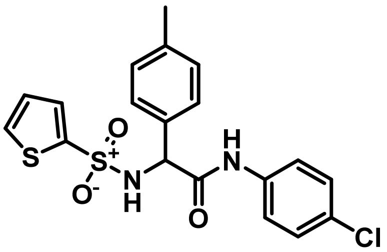


**N-(4-Chlorophenyl)-2-(thiophene-2-sulfonamido)-2-(p-tolyl)acetamide (Table 3; Analog 37):** This compound was prepared from **Method 4** using 4-chloroaniline and 2-(4-fluorophenyl)-2-(thiophene-2-sulfonamido)acetic acid (This compound was prepared from **Method 3**) to afford N-(4-chlorophenyl)-2-(4-fluorophenyl)-2-(thiophene-2-sulfonamido)acetamide as a TFA salt. LCMS: *m/z* $(M+H)^+$ = 421; $^1$H NMR (400 MHz, dmso) δ 10.35 (s, 1H), 8.94 (d, *J* = 9.3 Hz, 1H), 7.78 (dd, *J* = 5.0, 1.4 Hz, 1H), 7.49 – 7.40 (m, 2H), 7.35 – 7.23 (m, 3H), 7.08 (d, *J* = 7.9 Hz, 2H), 6.99 (dd, *J* = 5.0, 3.7 Hz, 1H), 5.14 (d, *J* = 9.3 Hz, 1H), 2.22 (s, 3H).

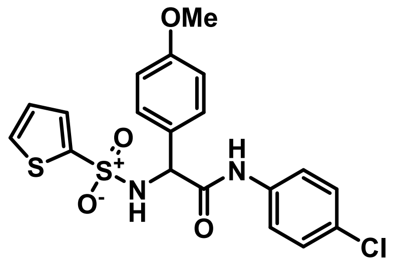


**N-(4-Chlorophenyl)-2-(4-methoxyphenyl)-2-(thiophene-2-sulfonamido)acetamide (Table 3; Analog 38):** This compound was prepared from **Method 4** using 4-chloroaniline and 2-(4-fluorophenyl)-2-(thiophene-2-sulfonamido)acetic acid (This compound was prepared from **Method 3**) to afford N-(4-chlorophenyl)-2-(4-fluorophenyl)-2-(thiophene-2-sulfonamido)acetamide as a TFA salt. LCMS: *m/z* $(M+H)^+$ = 437; $^1H$ NMR (400 MHz, dmso) δ 10.33 (s, 1H), 8.91 (d, *J* = 9.3 Hz, 1H), 7.78 (dd, *J* = 5.0, 1.4 Hz, 1H), 7.48 – 7.40 (m, 3H), 7.35 – 7.26 (m, 3H), 6.99 (dd, *J* = 5.0, 3.7 Hz, 1H), 6.87 – 6.79 (m, 2H), 5.12 (d, *J* = 9.3 Hz, 1H), 3.69 (s, 3H).

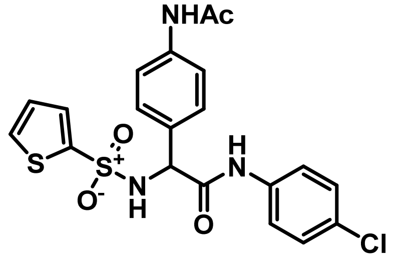


**2-(4-Acetamidophenyl)-N-(4-chlorophenyl)-2-(thiophene-2-sulfonamido)acetamide (Table 3; Analog 39):** This compound was prepared from **Method 4** using 4-chloroaniline and 2-(4-fluorophenyl)-2-(thiophene-2-sulfonamido)acetic acid (This compound was prepared from **Method 3**) to afford N-(4-chlorophenyl)-2-(4-fluorophenyl)-2-(thiophene-2-sulfonamido)acetamide as a TFA salt. LCMS: *m/z* $(M+H)^+$ = 464; $^1H$ NMR (400 MHz, dmso) δ 10.35 (s, 1H), 9.91 (s, 1H), 8.93 (d, *J* = 9.3 Hz, 1H), 7.78 (dd, *J* = 5.0, 1.4 Hz, 1H), 7.49 – 7.41 (m, 5H), 7.34 – 7.25 (m, 4H), 6.99 (dd, *J* = 5.0, 3.7 Hz, 1H), 5.13 (d, *J* = 9.3 Hz, 1H), 1.99 (s, 3H).

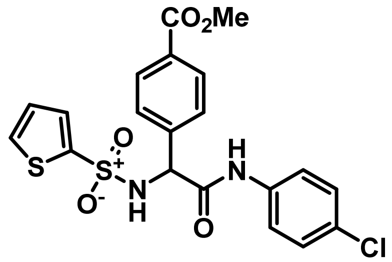

**Ethyl 4-(2-((4-chlorophenyl)amino)-2-oxo-1-(thiophene-2-sulfonamido)ethyl)benzoate (Table 3; Analog 40):** This compound was prepared from **Method 4** using 4-chloroaniline and 2-(4-fluorophenyl)-2-(thiophene-2-sulfonamido)acetic acid (This compound was prepared from **Method 3**) to afford N-(4-chlorophenyl)-2-(4-fluorophenyl)-2-(thiophene-2-sulfonamido)acetamide as a TFA salt. LCMS: *m/z* $(M+H)^+$ = 465; $^1$H NMR (400 MHz, dmso) δ 10.50 (s, 1H), 9.16 (d, *J* = 9.6 Hz, 1H), 7.90 – 7.83 (m, 2H), 7.77 (dd, *J* = 5.0, 1.4 Hz, 1H), 7.53 (d, *J* = 8.2 Hz, 2H), 7.49 – 7.40 (m, 3H), 7.36 – 7.28 (m, 2H), 6.97 (dd, *J* = 5.0, 3.7 Hz, 1H), 5.31 (d, *J* = 9.5 Hz, 1H), 3.81 (s, 3H).

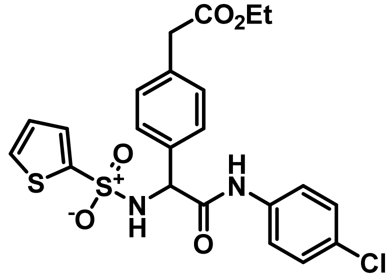

**Ethyl 2-(4-(2-((4-chlorophenyl)amino)-2-oxo-1-(thiophene-2-sulfonamido)ethyl)phenyl)acetate (Table 3; Analog 41):** This compound was prepared from **Method 4** using 4-chloroaniline and 2-(4-(2-ethoxy-2-oxoethyl)phenyl)-2-(thiophene-2-sulfonamido)acetic acid (This compound was prepared from **Method 3**) to afford ethyl 2-(4-(2-((4-chlorophenyl)amino)-2-oxo-1-(thiophene-2-sulfonamido)ethyl)phenyl)acetate as a TFA salt. LCMS: *m/z* $(M+H)^+$ = 493; $^1$H NMR (400 MHz, dmso) δ 10.39 (s, 1H), 8.99 (d, *J* = 9.4 Hz, 1H), 7.77 (dd, *J* = 5.0, 1.4 Hz, 1H), 7.49 – 7.40 (m, 3H), 7.36 – 7.27 (m, 4H), 7.18 (d, *J* = 8.0 Hz, 2H), 6.99 (dd, *J* = 5.0, 3.7 Hz, 1H), 5.18 (d, *J* = 9.3 Hz, 1H), 4.04 (q, *J* = 7.1 Hz, 2H), 3.59 (s, 2H), 1.15 (t, *J* = 7.1 Hz, 3H).

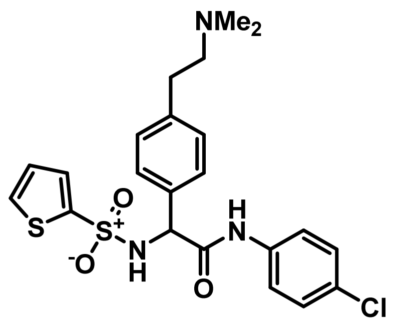

**N-(4-Chlorophenyl)-2-(4-(2-(dimethylamino)ethyl)phenyl)-2-(thiophene-2-sulfonamido)acetamide (Table 3; Analog 42):** This compound was prepared from **Method 4** using 4-chloroaniline and 2-(4-(2-(dimethylamino)ethyl)phenyl)-2-(thiophene-2-sulfonamido)acetic acid (This compound was prepared from **Method 3**) to afford N-(4-chlorophenyl)-2-(4-(2-(dimethylamino)ethyl)phenyl)-2-(thiophene-2-sulfonamido)acetamide as a TFA salt. LCMS: *m/z* $(M+H)^+$ = 478; $^1$H NMR (400 MHz, dmso) δ 10.41 (s, 1H), 9.42 (s, 1H), 9.00 (d, *J* = 9.5 Hz, 1H), 7.85 – 7.73 (m, 1H), 7.49 – 7.40 (m, 3H), 7.38 (d, *J* = 8.0 Hz, 2H), 7.34 – 7.27 (m, 2H), 7.20 (d, *J* = 8.0 Hz, 2H), 6.99 (dd, *J* = 5.0, 3.7 Hz, 1H), 5.19 (d, *J* = 9.4 Hz, 1H), 3.26 – 3.13 (m, 2H), 3.08 (d, *J* = 7.6 Hz, 1H), 2.89 (q, *J* = 9.2 Hz, 2H), 2.80 (d, *J* = 5.1 Hz, 2H), 2.78 (s, 6H).

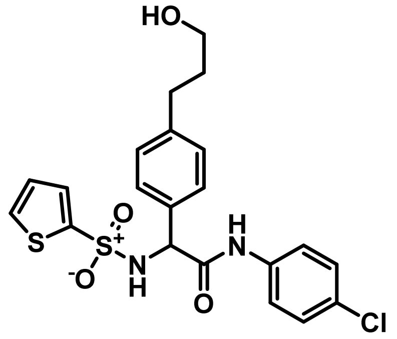

**N-(4-Chlorophenyl)-2-(4-(3-hydroxypropyl)phenyl)-2-(thiophene-2-sulfonamido)acetamide (Table 3; Analog 43):** This compound was prepared from **Method 4** using 4-chloroaniline and 2-(4-(3-hydroxypropyl)phenyl)-2-(thiophene-2-sulfonamido)acetic acid (This compound was prepared from **Method 3**) to afford N-(4-chlorophenyl)-2-(4-(3-hydroxypropyl)phenyl)-2-(thiophene-2-sulfonamido)acetamide as a TFA salt. LCMS: *m/z* $(M+H)^+$ = 465; $^1$H NMR (400 MHz, dmso) δ 10.36 (s, 1H), 8.93 (d, *J* = 9.4 Hz, 1H), 7.76 (dd, *J* = 5.0, 1.4 Hz, 1H), 7.49 – 7.41 (m, 2H), 7.36 – 7.25 (m, 4H), 7.09 (d, *J* = 8.1 Hz, 2H), 6.98 (dd, *J* = 5.0, 3.7 Hz, 1H), 5.15 (d, *J* =

9.3 Hz, 1H), 4.43 (t, $J$ = 5.1 Hz, 1H), 3.08 (qd, $J$ = 7.2, 4.3 Hz, 2H), 2.56 – 2.48 (m, 2H), 1.69 – 1.58 (m, 2H).

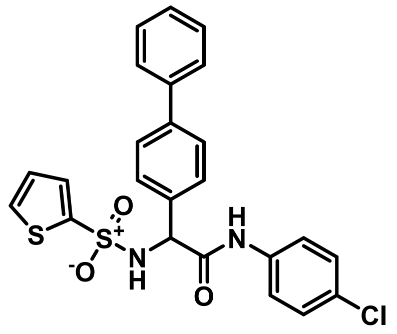


**2-([1,1'-Biphenyl]-4-yl)-N-(4-chlorophenyl)-2-(thiophene-2-sulfonamido)acetamide (Table 3; Analog 44):** This compound was prepared from **Method 4** using 4-chloroaniline and 2-([1,1'-biphenyl]-4-yl)-2-(thiophene-2-sulfonamido)acetic acid (This compound was prepared from **Method 3**) to afford 2-([1,1'-biphenyl]-4-yl)-N-(4-chlorophenyl)-2-(thiophene-2-sulfonamido)acetamide as a TFA salt. LCMS: *m/z* $(M+H)^+$ = 483; $^1H$ NMR (400 MHz, dmso) δ 10.45 (s, 1H), 9.06 (d, $J$ = 9.3 Hz, 1H), 7.78 (dd, $J$ = 5.0, 1.4 Hz, 1H), 7.59 (ddd, $J$ = 11.0, 7.8, 1.7 Hz, 4H), 7.52 – 7.39 (m, 7H), 7.38 – 7.28 (m, 3H), 6.99 (dd, $J$ = 5.0, 3.7 Hz, 1H), 5.25 (d, $J$ = 9.4 Hz, 1H).

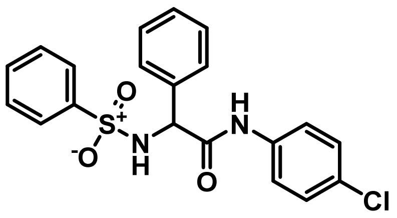


**N-(4-Chlorophenyl)-2-phenyl-2-(phenylsulfonamido)acetamide (Table 4; Analog 45):** This compound was prepared from **Method 4** using 4-chloroaniline and 2-phenyl-2-(phenylsulfonamido)acetic acid (This compound was prepared from **Method 3**) to afford N-(4-chlorophenyl)-2-phenyl-2-(phenylsulfonamido)acetamide as a TFA salt. LCMS: *m/z* $(M+H)^+$ = 401; $^1H$ NMR (400 MHz, dmso) δ 10.36 (s, 1H), 8.78 (d, $J$ = 9.5 Hz, 1H), 7.76 – 7.68 (m, 2H), 7.50 – 7.16 (m, 11H), 5.14 (d, $J$ = 9.5 Hz, 1H).

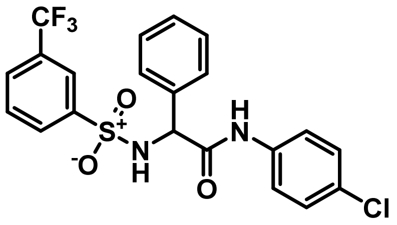

**N-(4-Chlorophenyl)-2-phenyl-2-((3-(trifluoromethyl)phenyl)sulfonamido)acetamide (Table 4; Analog 46):** This compound was prepared from **Method 4** using 4-chloroaniline and 2-phenyl-2-((3-(trifluoromethyl)phenyl)sulfonamido)acetic acid (This compound was prepared from **Method 3**) to afford N-(4-chlorophenyl)-2-phenyl-2-((3-(trifluoromethyl)phenyl)-sulfonamido)acetamide as a TFA salt. LCMS: *m/z* $(M+H)^+$ = 469; $^1$H NMR (400 MHz, cdcl3) δ 7.87 – 7.79 (m, 2H), 7.58 (d, *J* = 7.8 Hz, 1H), 7.40 (t, *J* = 7.8 Hz, 1H), 7.32 – 7.25 (m, 2H), 7.19 – 7.08 (m, 7H), 5.15 (d, *J* = 9.3 Hz, 1H).

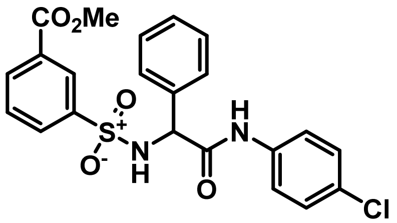

**Methyl 3-(N-(2-((4-chlorophenyl)amino)-2-oxo-1-phenylethyl)sulfamoyl)benzoate (Table 4; Analog 47):** This compound was prepared from **Method 4** using 4-chloroaniline and 2-phenyl-2-((3-(trifluoromethyl)phenyl)sulfonamido)acetic acid (This compound was prepared from **Method 3**) to afford N-(4-chlorophenyl)-2-phenyl-2-((3-(trifluoromethyl)phenyl)-sulfonamido)acetamide as a TFA salt. LCMS: *m/z* $(M+H)^+$ = 459; $^1$H NMR (400 MHz, cdcl3) δ 8.32 (t, *J* = 1.8 Hz, 1H), 8.11 (dt, *J* = 7.8, 1.4 Hz, 1H), 7.87 (dt, *J* = 8.0, 1.5 Hz, 1H), 7.50 – 7.41 (m, 2H), 7.32 – 7.15 (m, 8H), 6.09 (d, *J* = 5.5 Hz, 1H), 5.15 (d, *J* = 9.3 Hz, 1H), 3.91 (s, 3H).

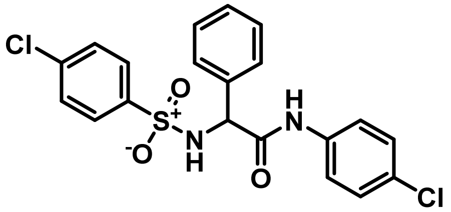

**N-(4-Chlorophenyl)-2-((4-chlorophenyl)sulfonamido)-2-phenylacetamide (Table 4; Analog 48):** This compound was prepared from **Method 4** using 4-chloroaniline and 2-((4-

chlorophenyl)sulfonamido)-2-phenylacetic acid (This compound was prepared from **Method 3**) to afford N-(4-chlorophenyl)-2-((4-chlorophenyl)sulfonamido)-2-phenylacetamide as a TFA salt. LCMS: *m/z* $(M+H)^+$ = 435; $^1$H NMR (400 MHz, cdcl3) δ 7.61 – 7.53 (m, 2H), 7.31 – 7.17 (m, 5H), 7.17 (m, 5H), 7.15 (d, *J* = 2.3 Hz, 1H), 5.15 (d, *J* = 9.3 Hz, 1H).

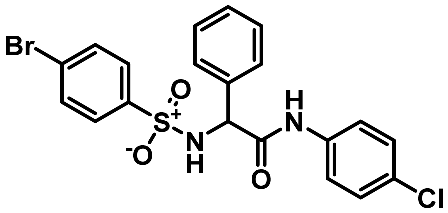


**2-((4-Bromophenyl)sulfonamido)-N-(4-chlorophenyl)-2-phenylacetamide (Table 4; Analog 49):** This compound was prepared from **Method 4** using 4-chloroaniline and 2-((4-bromophenyl)sulfonamido)-2-phenylacetic acid (This compound was prepared from **Method 3**) to afford 2-((4-bromophenyl)sulfonamido)-N-(4-chlorophenyl)-2-phenylacetamide as a TFA salt. LCMS: *m/z* $(M+H)^+$ = 481; $^1$H NMR (400 MHz, cdcl3) δ 7.52 – 7.44 (m, 2H), 7.41 – 7.33 (m, 2H), 7.28 – 7.20 (m, 4H), 7.17 – 7.10 (m, 5H), 4.95 (d, *J* = 1.5 Hz, 1H).

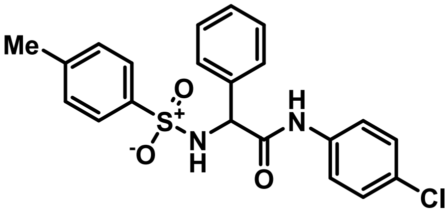


**N-(4-Chlorophenyl)-2-((4-methylphenyl)sulfonamido)-2-phenylacetamide (Table 4; Analog 50):** This compound was prepared from **Method 4** using 4-chloroaniline and 2-((4-methylphenyl)sulfonamido)-2-phenylacetic acid (This compound was prepared from **Method 3**) to afford N-(4-chlorophenyl)-2-((4-methylphenyl)sulfonamido)-2-phenylacetamide as a TFA salt. LCMS: *m/z* $(M+H)^+$ = 415; $^1$H NMR (400 MHz, cdcl3) δ 7.58 – 7.51 (m, 1H), 7.30 – 7.20 (m, 2H), 7.20 – 7.03 (m, 6H), 4.89 (d, *J* = 1.5 Hz, 1H), 2.21 (s, 3H).

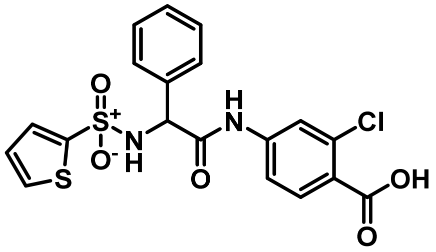

**2-Chloro-4-(2-phenyl-2-(thiophene-2-sulfonamido)acetamido)benzoic acid (Table 5; Analog 53):** This compound was prepared from **Method 4** using methyl 4-amino-2-chlorobenzoate to afford methyl 2-chloro-4-(2-phenyl-2-(thiophene-2-sulfonamido)acetamido)benzoate. Methyl 2-chloro-4-(2-phenyl-2-(thiophene-2-sulfonamido)acetamido)benzoate in MeOH was added 4M NaOH and stirred for 12hrs at 50ºC to aford 2-chloro-4-(2-phenyl-2-(thiophene-2-sulfonamido)acetamido)benzoic acid as a TFA salt. LCMS: *m/z* $(M+H)^+$ = 451; $^1$H NMR (400 MHz, cdcl$_3$) δ 8.84 (s, 1H), 7.49 – 7.41 (m, 3H), 7.35 – 7.25 (m, 4H), 6.97 (t, *J* = 7.7 Hz, 1H), 6.86 (d, *J* = 8.3 Hz, 1H), 6.31 (d, *J* = 7.2 Hz, 1H), 5.40 (q, *J* = 5.9 Hz, 1H).

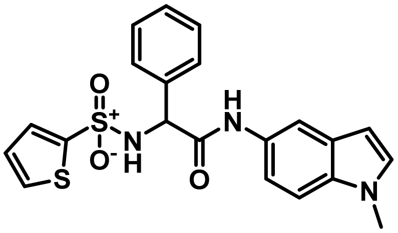

**2-Chloro-4-(2-phenyl-2-(thiophene-2-sulfonamido)acetamido)benzoic acid (Table 5; Analog 54):** This compound was prepared from **Method 4** using 1-methyl-1H-indol-5-amine to afford N-(1-methyl-1H-indol-5-yl)-2-phenyl-2-(thiophene-2-sulfonamido)acetamide as a TFA salt. LCMS: *m/z* $(M+H)^+$ = 426; $^1$H NMR (400 MHz, cdcl$_3$) δ 7.63 (d, *J* = 2.0 Hz, 1H), 7.50 (ddd, *J* = 13.1, 4.4, 1.3 Hz, 2H), 7.33-7.32 (m, 3H), 7.30 (s, 1H), 7.21 (d, *J* = 8.7 Hz, 1H), 7.18 – 7.04 (m, 1H), 7.08 – 7.01 (m, 1H), 6.97 (dd, *J* = 5.0, 3.7 Hz, 1H), 6.41 (d, *J* = 3.1 Hz, 1H), 6.16 (d, *J* = 5.0 Hz, 1H), 5.02 (d, *J* = 5.0 Hz, 1H), 3.76 (s, 3H).

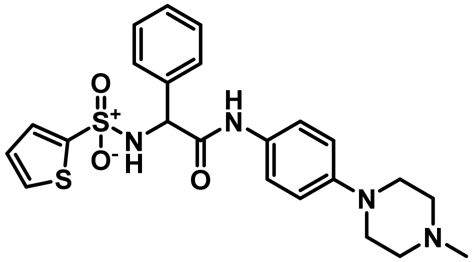

**N-(4-(4-Methylpiperazin-1-yl)phenyl)-2-phenyl-2-(thiophene-2-sulfonamido)acetamide (Table 5; Analog 55):** This compound was prepared from **Method 4** using 4-(4-methylpiperazin-1-yl)aniline to afford N-(4-(4-methylpiperazin-1-yl)phenyl)-2-phenyl-2-(thiophene-2-sulfonamido)acetamide as a TFA salt. LCMS: *m/z* $(M+H)^+$ = 471; $^1$H NMR (400 MHz, dmso) δ 10.41 (s, 1H), 9.01 (d, *J* = 9.4 Hz, 1H), 7.77 (dd, *J* = 5.0, 1.4 Hz, 1H), 7.49 – 7.20 (m, 9H), 6.98 (dd, *J* = 5.0, 3.7 Hz, 1H), 5.20 (d, *J* = 9.4 Hz, 1H), 3.47-3.44(m, 4H), 2.39-2.35(m, 4H), 2.22(s, 3H).

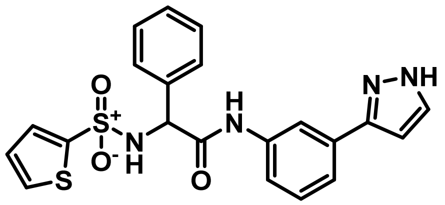


**N-(3-(1H-pyrazol-3-yl)phenyl)-2-phenyl-2-(thiophene-2-sulfonamido)acetamide (Table 5; Analog 56):** This compound was prepared from **Method 4** using 3-(1H-pyrazol-3-yl)aniline to afford N-(3-(1H-pyrazol-3-yl)phenyl)-2-phenyl-2-(thiophene-2-sulfonamido)acetamide as a TFA salt. LCMS: *m/z* $(M+H)^+$ = 470.1; $^1$H NMR (400 MHz, DMSO-$d_6$) δ 8.58 (s, 1H), 8.19 (s, 1H), 8.10 (d, *J* = 8.6 Hz, 2H), 7.56 -7.47 (m, 2H), 7.49 – 7.41 (m, 1H), 7.35 (s, 1H), 7.33 (s, 1H), 3.01 (s, 3H), 2.96 (s, 3H), 2.29 (dq, *J* = 15.2, 7.6 Hz, 2H), 2.16 (dq, *J* = 15.0, 7.5 Hz, 2H), 1.84 (s, 3H), 1.06 (t, J = 7.6 Hz, 6H).

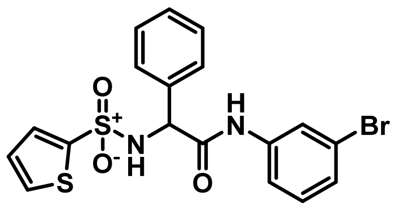


**N-(3-Bromophenyl)-2-phenyl-2-(thiophene-2-sulfonamido)acetamide (Table 5; Analog 57):** This compound was prepared from **Method 4** using 3-(1H-pyrazol-3-yl)aniline to afford N-(3-bromophenyl)-2-phenyl-2-(thiophene-2-sulfonamido)acetamide as a TFA salt. LCMS: *m/z* $(M+H)^+$ = 452; $^1$H NMR (400 MHz, dmso) δ 10.40 (s, 1H), 9.0 (d, *J* = 9.4 Hz, 1H), 7.5 (dd, *J* = 5.0, 1.4 Hz, 1H), 7.47 – 7.18 (m, 8H), 6.96 (dd, *J* = 5.0, 3.7 Hz, 1H), 5.20 (d, *J* = 9.4 Hz, 1H).

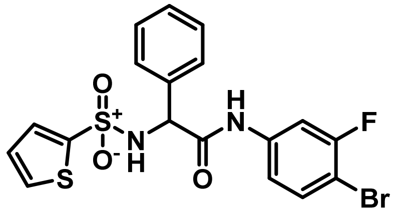


**N-(4-Bromo-3-fluorophenyl)-2-phenyl-2-(thiophene-2-sulfonamido)acetamide (Table 5; Analog 58):** This compound was prepared from **Method 4** using 4-bromo-3-fluoroaniline to afford N-(4-bromo-3-fluorophenyl)-2-phenyl-2-(thiophene-2-sulfonamido)acetamide as a TFA salt. LCMS: *m/z* $(M+H)^+$ = 469; $^1$H NMR (400 MHz, dmso) δ 10.40 (s, 1H), 8.99 (d, *J* = 9.1 Hz, 1H), 7.81 – 7.68 (m, 1H), 7.53 – 7.46 (m, 2H), 7.50 – 7.39 (m, 2H), 7.42 – 7.27 (m, 2H), 7.18 – 7.05 (m, 2H), 6.99 (dd, *J* = 5.0, 3.7 Hz, 1H), 5.44 (d, *J* = 9.1 Hz, 1H).

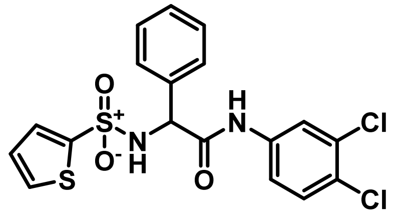


**N-(3,4-Dichlorophenyl)-2-phenyl-2-(thiophene-2-sulfonamido)acetamide (Table 5; Analog 59):** This compound was prepared from **Method 4** using 3,4-dichloroaniline to afford N-(3,4-dichlorophenyl)-2-phenyl-2-(thiophene-2-sulfonamido)acetamide as a TFA salt. LCMS: *m/z* $(M+H)^+$ = 443; $^1$H NMR (400 MHz, dmso) δ 10.44 (s, 1H), 9.09 (d, *J* = 9.5 Hz, 1H), 7.78 (dd, *J* = 5.0, 1.4 Hz, 1H), 7.48 – 7.28 (m, 8H), 6.99 (dd, *J* = 5.0, 3.7 Hz, 1H), 5.22 (d, *J* = 9.4 Hz, 1H).

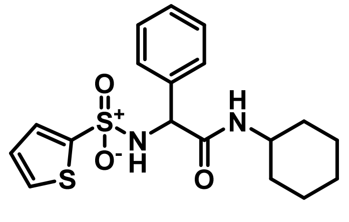


**N-Cyclohexyl-2-phenyl-2-(thiophene-2-sulfonamido)acetamide (Table 5; Analog 60):** This compound was prepared from **Method 4** using cyclohexanamine to afford N-cyclohexyl-2-phenyl-2-(thiophene-2-sulfonamido)acetamide as a TFA salt. LCMS: *m/z* $(M+H)^+$ = 379; $^1$H NMR (400 MHz, dmso) δ 8.72 (d, *J* = 9.7 Hz, 1H), 8.03 (d, *J* = 7.7 Hz, 1H), 7.81 (d, *J* = 5.2 Hz, 1H), 7.46 – 7.41 (m, 1H), 7.32 (d, *J* = 7.4 Hz, 2H), 7.29 – 7.16 (m, 2H), 7.04 (t, *J* = 4.4 Hz, 1H),

5.01 (d, $J$ = 9.7 Hz, 1H), 1.60 (d, $J$ = 12.4 Hz, 2H), 1.46 (d, $J$ = 12.6 Hz, 2H), 1.22 – 0.90 (m, 6H).

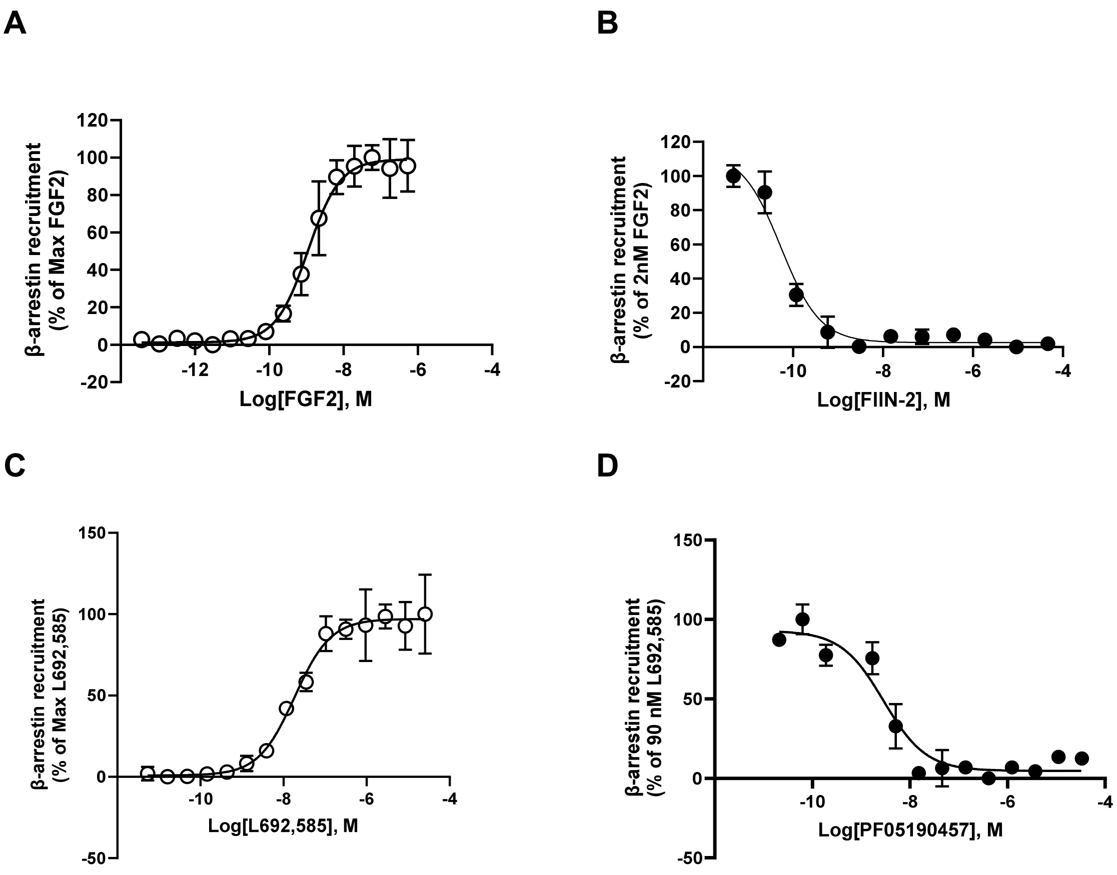


**Supplemental figure 1.** Performance of counter assays. **A.** Dose response curve of PathHunter FGFR1 functional assay in agonist mode. $EC_{50}$ of FGF2, an endogenous ligand of FGFR1 receptor, was 1.1nM. **B.** Dose response curve of PathHunter FGFR1 functional assay in antagonist mode. ~$EC_{80}$ FGF2 (2 nM) was used to stimulate receptors. $IC_{50}$ of FIIN-2, an antagonist of FGFR1 receptor, was 0.071 nM. **C.** Dose response curve of PathHunter GHSR1a β-arrestin assay in agonist mode. $EC_{50}$ of L692,585, an agonist of GHSR1a receptor, was 18.4nM. **D.** Dose response curve of PathHunter GHSR1a β-arrestin assay in antagonist mode. ~$EC_{80}$ L692,585 (90 nM) was used to stimulate receptors. $IC_{50}$ of PF05190457, an antagonist of FGFR1 receptor, was 2.9 nM.

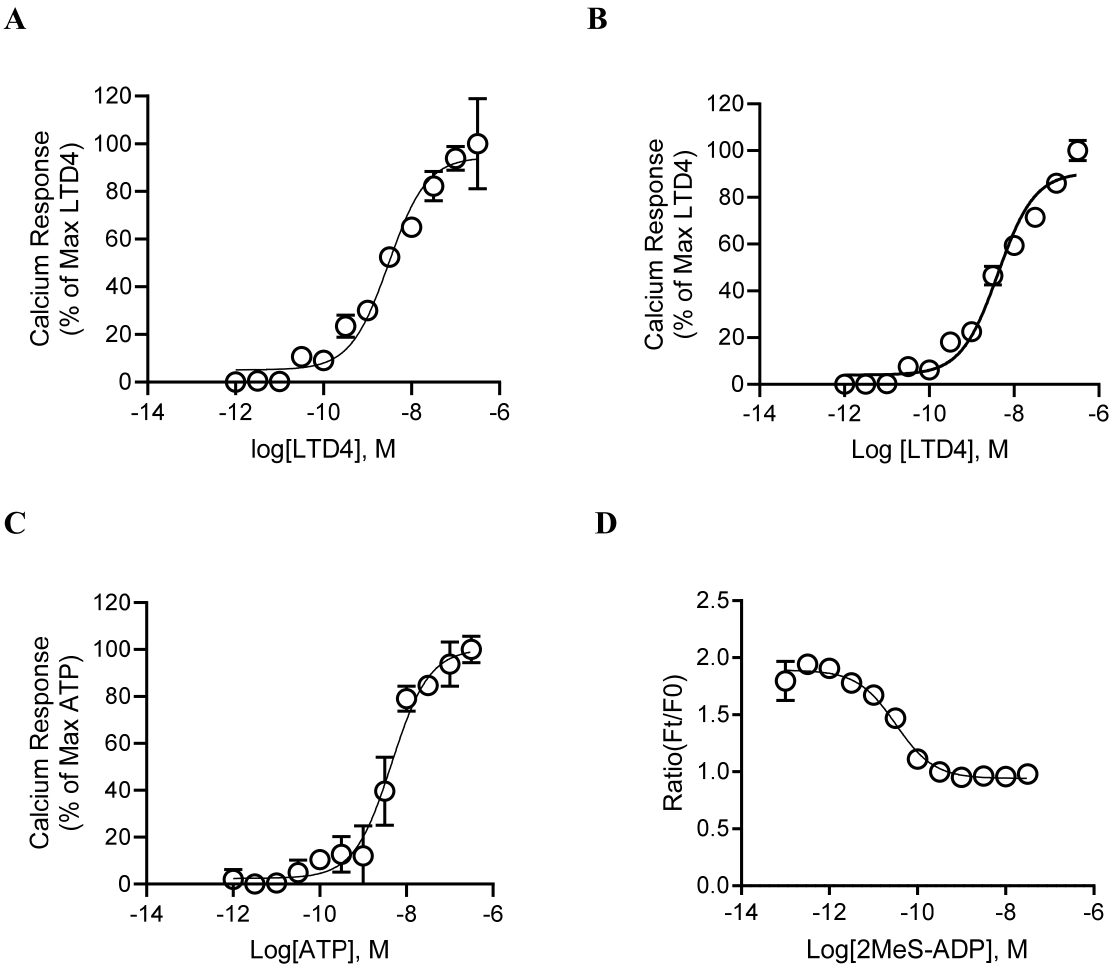


**Supplemental figure 2.** Performance of selectivity assays. **A**. Dose-response curve of LTD4 (EC50=3.0 nM) in calcium mobilization assay in HEK293 hCysLTR1 stable cell line. **B**. Dose-response curve of LTD4 (EC50=4.2 nM) in calcium mobilization assay in HEK293 hCysLTR2 stable cell line. **C**. Dose-response curve of ATP (EC50=4.6 nM) in calcium mobilization assay in HEK293 hP2RY1 stable cell line. **D**. Dose-response curve of LTD4 (EC50=0.032 nM) in ACTone cAMP assay in HEK293 hP2RY12 stable cell line.

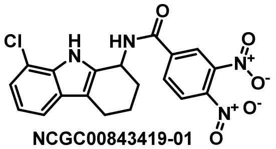
NCGC00843419-01

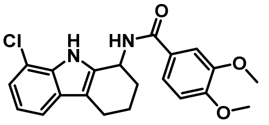
NCGC00423978-04

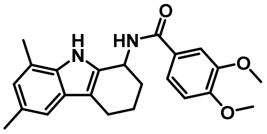
NCGC00843850-01

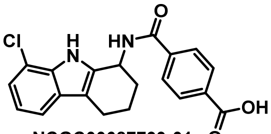
NCGC00687793-01

| Compound ID | $IC_{50}$ (μM) | Efficacy (%) |
|---|---|---|
| NCGC00843419-01 | 12.65 | -74 |
| NCGC00423978-04 | 2.7 | -111 |
| NCGC00843850-01 | 11.27 | -87 |
| NCGC00687793-01 | 28.31 | -49 |

| Compound ID | $IC_{50}$ (μM) | Efficacy (%) |
|---|---|---|
| NCGC00105527-02 | 6 | -108 |
| NCGC00645518-01 | 13.43 | -50 |
| NCGC00689480-01 | 11.27 | -111 |

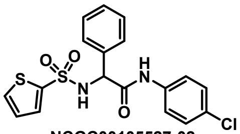
NCGC00105527-02

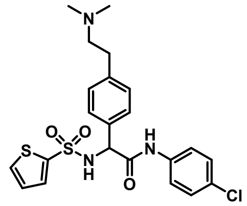
NCGC00645518-01

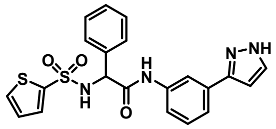
NCGC00689480-01

**Supplemental figure 3.** Table showing chemical structure and activity of analogs tested in Figure 7 in the calcium mobilization assay.

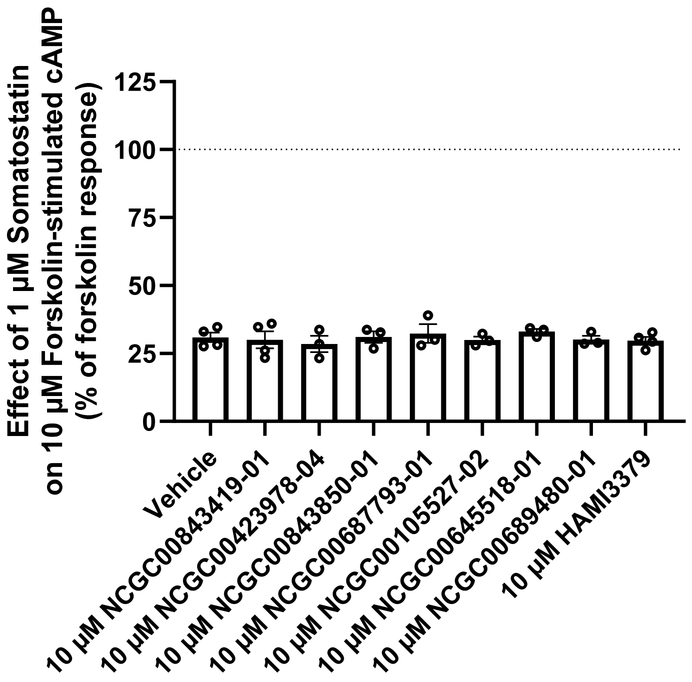


**Supplemental figure 4**. Assessment of off-target effects on cAMP signaling in GLUTag cells. GLUTag cells were transduced with a control adenovirus and the effects of test compounds on 1 µM somatostatin-mediated inhibition of 10 µM forskolin-stimulated cAMP were measured. Data were normalized to the forskolin response within each test compound treatment condition and represent mean ± SEM of three to four independent experiments performed in triplicate wells. Data were analyzed by unpaired t-test.

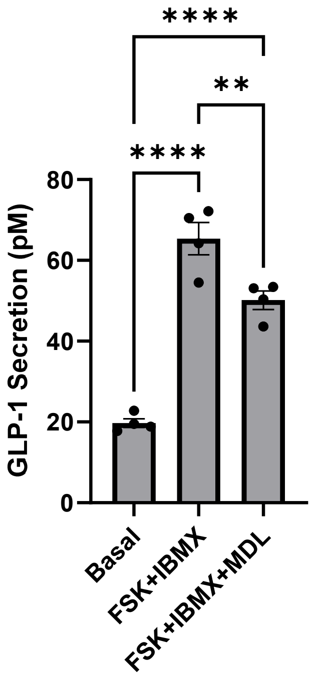


**Supplemental figure 5**. GPR17-mediated inhibition of GLP-1 secretion in GLUTag cells. GLUTag cells were transduced with adenovirus containing hGPR17L. Stimulating hGPR17L expressing GLUTag cells with 10 µM forskolin and 10 µM IBMX caused increase of GLP-1 secretion. Activation of hGPR17L with 6.3 µM MDL inhibited the forskolin and IBMX-stimulated GLP-1 secretion. Data were analyzed by one-way ANOVA and then followed by Tukey's multiple comparison test.

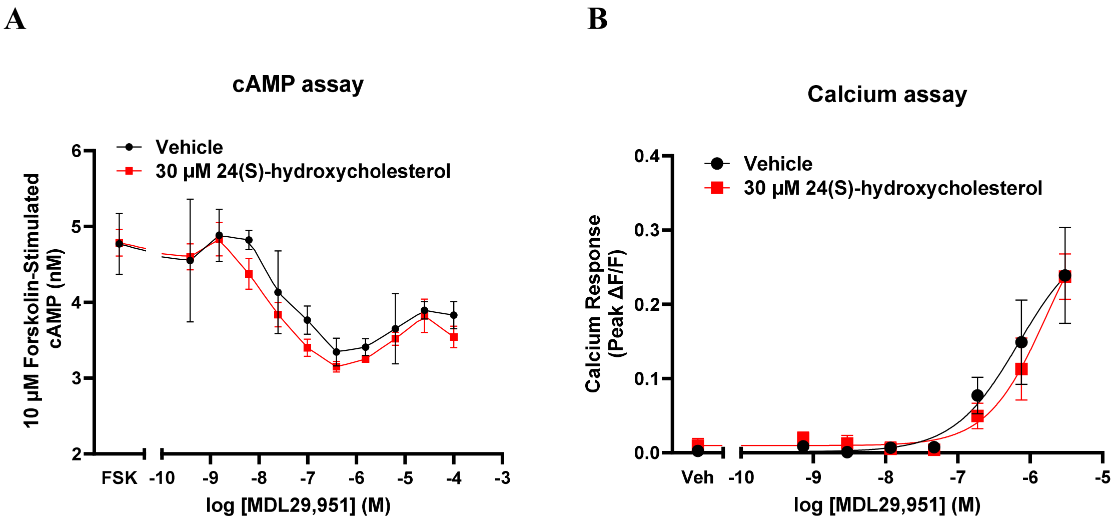


**Supplemental figure 6.** Assessment of modulation of 24(S)-hydroxycholesterol on GPR17. GLUTag cells were transduced with adenovirus containing hGPR17L. Simulation of GPR17 by 24(S)-hydroxycholesterol was evaluated under the condition without MDL, or with increase concentration of MDL in both cAMP accumulation assay (**A**) and calcium mobilization assay (**B**).
